# Wall stiffening is a primary contributor to motility loss in crohn’s disease: an electromechanical modeling study

**DOI:** 10.64898/2026.09.20.752998

**Authors:** Ahilan Anantha Krishnan, Masah Abubaker, Maria A. Holland

## Abstract

Fibrotic strictures are among the most disabling complications of Crohn’s disease, permanently narrowing the bowel and impairing motility, yet no approved therapy reverses them. Chronic inflammation alters pacemaker-network coupling, smooth-muscle excitability, and calcium-dependent contractility, while fibrosis thickens the bowel wall, narrows the lumen, and changes tissue mechanics. The relative contributions of these coupled electrical, contractile, and structural alterations to motility loss remain unclear. To address this gap, we develop an integrated electromechanical finite-element framework for fibrostenosing Crohn’s disease that couples a fibrosis-driven growth model with a FitzHugh-Nagumo electromechanical model. A full-factorial 2^5^ design of experiments is used to quantify the relative effects of electrical diffusivity, excitation threshold, peak active stress, wall stiffness, and hypertrophic remodeling on cyclic lumen-volume deformation. Motility is quantified by the standard deviation of lumen volume over one contraction cycle. Within the parameter ranges examined, increased wall stiffness emerged as the dominant contributor to motility loss, followed by impaired smooth-muscle contractility. Changes in excitation threshold, hypertrophic remodeling, and electrical diffusivity produced substantially smaller effects. Pairwise interactions were small relative to the dominant main effects, indicating that the mechanisms contributed largely through their individual effects. Our findings suggest that limiting wall stiffening while preserving smoothmuscle contractile function may provide a therapeutic strategy for maintaining intestinal motility in fibrostenosing Crohn’s disease.

## 1 Introduction

Inflammatory bowel disease (IBD) affects more than six million people worldwide, with Crohn’s disease accounting for approximately 40% of cases; Crohn’s disease is a chronic relapsing disorder whose incidence continues to rise across newly industrialized nations (Ng et al. 2017; Alatab et al. 2020). Among the most clinically debilitating complications of Crohn’s is the development of fibrotic strictures in the ileum, the terminal part of the small intestine. In these cases, progressive remodeling of the bowel wall permanently narrows the intestinal lumen, disrupts coordinated intestinal motility (Menys et al. 2018; Beek et al. 2025), and frequently leaves patients with no therapeutic option beyond endoscopic dilation or surgical resection. Strictures are fairly common; along with penetrating complications, they occur in approximately one-third of patients within ten years of diagnosis and in most patients over the disease lifetime (Cosnes et al. 2002; Rieder et al. 2013). Once established, fibrotic strictures are largely refractory to current anti-inflammatory therapies, and no approved treatment reverses the associated smooth muscle and extracellular matrix remodeling (Rieder et al. 2017; Bettenworth et al. 2024).

Depletion of interstitial cells of Cajal (ICC), which form the pacemaker network responsible for generating the electrical slow waves that coordinate intestinal motility, has been documented in Crohn’s disease small-intestinal tissue (Porcher et al. 2002; Wang et al. 2007). Functional studies further demonstrate altered smooth muscle function in IBD, including disrupted calcium signalling and impaired circular muscle contraction (Johnson et al. 2023; Cao et al. 2005). Recent histological studies have further improved our understanding of Crohn’s disease, demonstrating that submucosal smooth muscle hyperplasia and muscularis propria hypertrophy are the dominant tissue alterations, producing wall thickening and luminal narrowing that exceed the contribution of collagen deposition (Chen et al. 2017; Veisman et al. 2024). Together, these findings suggest that chronic inflammation and the fibrosis it drives impair intestinal function through coupled changes in electrical conduction, active contraction, tissue mechanics, and wall geometry. However, the relative contributions of these factors to motility loss remain unresolved.

Chronic intestinal inflammation and the resulting fibrosis introduce several tightly coupled alterations that may contribute to motility loss. ICC depletion, driven primarily by the inflammatory response with fibrosis acting as an additional compounding factor, is expected to reduce slow-wave conduction velocity and electrical coupling through loss of the intestinal pacemaker network (Porcher et al. 2002; Wang et al. 2007). Partial depolarization of the smooth-muscle resting membrane potential has been observed in experimental colitis and is associated with reduced spike amplitude and altered excitability (Cohen et al. 1986). Active contractility is also impaired because inflammation disrupts the calcium signals needed for smooth muscle contraction. This occurs partly through reduced expression and impaired function of Cav1.2 L-type calcium channels (Liu et al. 2001; Shi and Sarna 2005; Ross et al. 2010). In parallel, inflammation-driven structural remodeling increases tissue stiffness and promotes progressive luminal narrowing, as observed in histological and elastographic measurements (Dal Buono et al. 2022; Lu et al. 2019; Chen et al. 2017). Determining how these mechanisms combine to produce motility loss requires a framework in which selected disease-associated parameters can be perturbed independently and in combination while the resulting functional consequences are quantified systematically. A full factorial design of experiments (DOE) therefore provides a natural strategy for decomposing these coupled effects into ranked main effects and pairwise interactions (Saltelli 2002; Brandstaeter et al. 2021). Similar approaches have successfully resolved dominant mechanisms in cardiac electrophysiology (Sobie 2009; Britton et al. 2013) and vascular wall mechanics (Heusinkveld et al. 2018), but have not yet been extended to intestinal motility or fibrostenosing disease.

Over the past two decades, gastrointestinal electrophysiology has been increasingly formalized through both phenomenological FitzHugh-Nagumo formulations and biophysically detailed ionic cell models, with applications ranging from one-dimensional representations of intestinal slow-wave propagation (Aliev et al. 2000) to quantitative descriptions of ICC and smooth-muscle cell (SMC) electrophysiology (Poh et al. 2012; Corrias and Buist 2007). More recently, fully coupled electromechanical finite element frameworks have demonstrated that wave propagation, active contraction, and luminal deformation can be captured within a unified continuum setting for both the stomach (Klemm et al. 2020; Klemm et al. 2023) and intestine (Djoumessi et al. 2024; Djoumessi et al. 2025). In parallel, growth and remodeling theory, based on the multiplicative decomposition of the deformation gradient (Rodriguez et al. 1994), enables irreversible wall thickening to be represented computationally and has been applied extensively to vascular (Bräu et al. 2017) and airway remodeling (Eskandari et al. 2015). Collectively, these advances now make it possible to study the consequences of Crohn’s disease-associated inflammation and the resulting fibrosis as an integrated electromechanical disease process rather than as isolated cellular or histological abnormalities. Yet, these modeling capabilities remain unutilized in Crohn’s disease. Without integrating inflammation-driven structural remodeling, altered electrophysiology, reduced active contraction, and altered tissue mechanics within a single framework, existing models cannot quantify their relative contributions to motility loss.

We therefore develop a two-step computational framework to investigate how fibrotic strictures alter ileal electromechanics and motility in Crohn’s disease. The framework couples inflammation-driven structural remodeling with intestinal electromechanics and is combined with a full-factorial 2^5^ DOE to quantify the relative contributions of five selected disease-associated mechanisms: reduced electrical diffusivity, altered excitation threshold, impaired active contractility, increased wall stiffness, and hypertrophic wall remodeling.

## 2 Methods

### 2.1 Model overview

Quantifying how inflammation- and fibrosis-associated alterations affect intestinal motility requires a model that captures growth, electrical conduction, active contraction, and structural wall mechanics within a single framework (Figure 1). It also requires independent perturbation of selected disease mechanisms so that their relative contributions can be quantified systematically. We therefore adopt a two-step sequential approach (Figure 2): Model 1 evolves a fibrosis field and drives irreversible wall growth. The resulting geometry and fibrosis distribution are transferred to Model 2, which simulates FitzHugh-Nagumo monodomain slow-wave propagation and active smooth-muscle contraction. A full-factorial 2^5^ design of experiments is used to compare five disease-associated factors. Hypertrophic remodeling, represented by *ϑ*^h^, determines the geometry generated by Model 1, whereas electrical diffusivity scaling *η*_*D*_, excitation-threshold scaling *η*_*a*_, active-tension scaling *η*_*T*_, and shear-modulus scaling *η*_*µ*_ modify the electromechanical response in Model 2. The resulting effects on the motility metric MM are used to rank the relative contributions of geometric remodeling, electrical alterations, impaired contractility, and wall stiffening. Both steps are implemented in Abaqus/Standard using user-defined material subroutines (UMAT/UMATHT), and the temperature degree of freedom is exploited as a surrogate scalar field to solve each reaction-diffusion equation (one for fibrosis-driven remodeling and another for electromechanics) within the existing structural solver (Table 1).

**Table 1.**
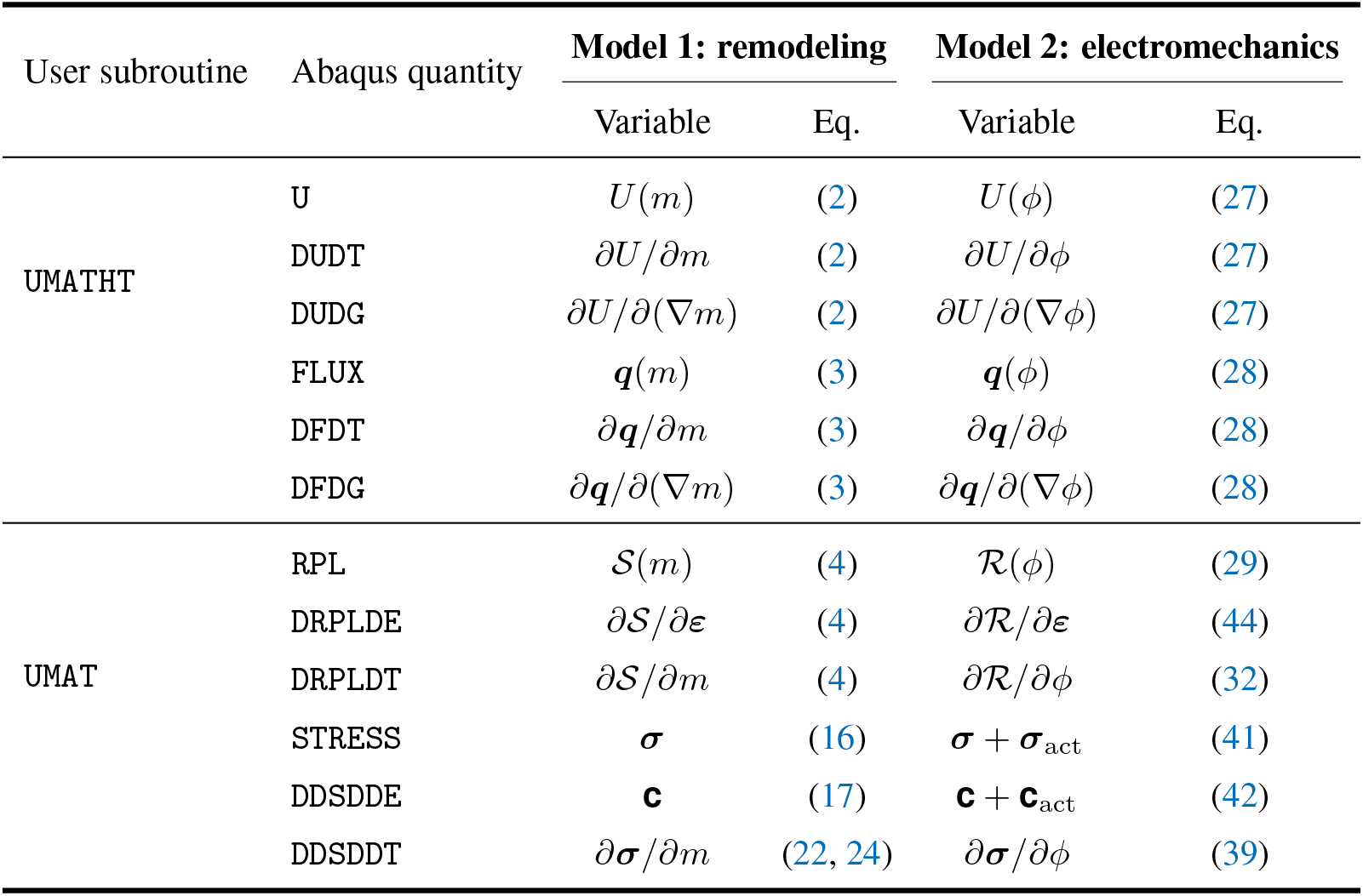
Terms used in Abaqus user subroutine for remodeling and electromechanical reaction-diffusion equations. The global Newton-Raphson tangent requires the internal energy (U) and flux via UMATHT and the reaction rate (RPL) and stress via UMAT, along with their derivatives.

**Figure 1.**
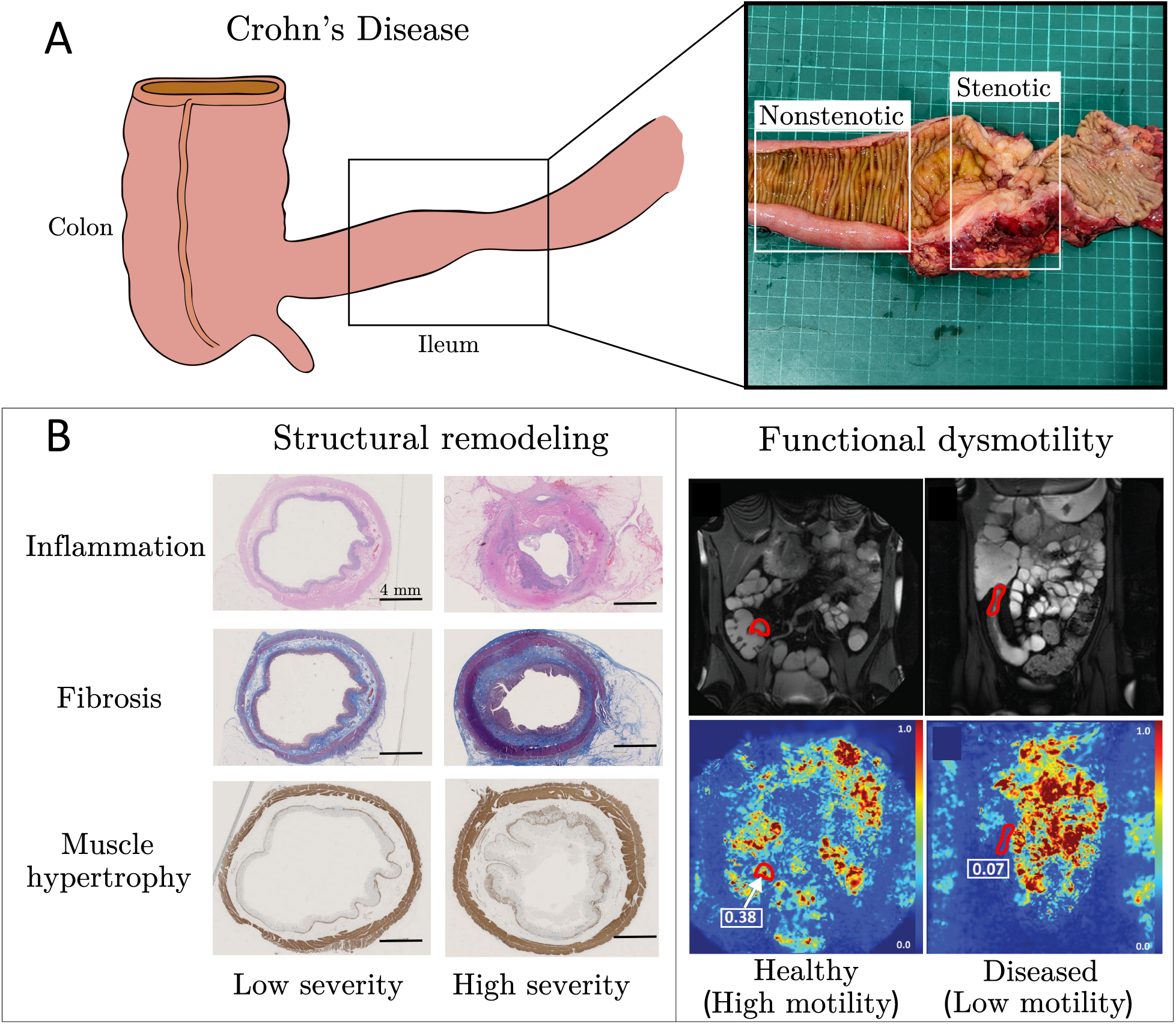
Pathological manifestations of Crohn’s disease and overview of the computational framework linking fibrotic remodeling to intestinal (ileum) dysmotility. **(A)** Schematic representation of the colon and ileum with gross pathology of a surgical specimen showing nonstenotic and stenotic ileal regions; fibrotic strictures cause permanent luminal narrowing that is refractory to antiinflammatory therapy (modified from Zhang et al. (2025) with permission). **(B)** Progressive structural and functional changes in Crohn’s disease bowel wall with increasing disease severity. **Left:** Representative histological cross-sections illustrating remodeling across acute and chronic inflammation (stained with haematoxylin and eosin), submucosal fibrosis (Masson’s trichrome), and muscularis propria hypertrophy (anti-desmin); progress from low to high indicates increasing severity (adapted from Zhao et al. (2021) under the Creative Commons license). **Right:** Magnetic Resonance (MR) images and motility maps showing changes from healthy (high-motility) to fibrotic (low-motility) terminal ileum, outlined in red (modified from Menys et al. (2018) with permission). The healthy subject exhibited high motility (0.38 arbitrary units (au)), reflecting normal bowel contractions. The diseased subject, with neoterminal ileal Crohn’s disease, exhibited low motility (0.07 au, which is below the 0.30 au cutoff for normal motility), demonstrating reduced bowel movement.

**Figure 2.**
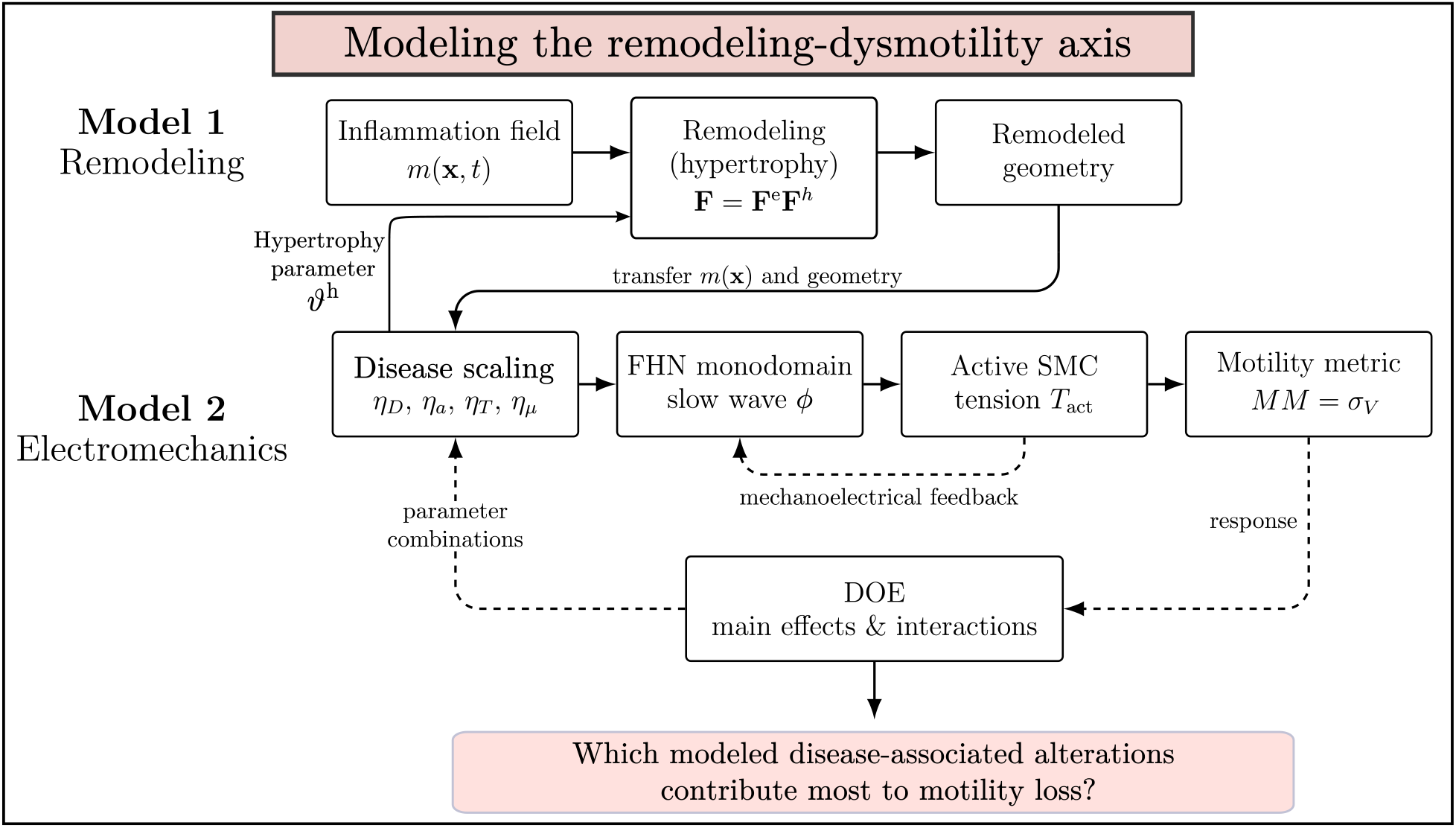
Schematic of the two-model computational framework. Model 1 captures fibrosis-driven structural remodeling through the fibrotic field *m*(***x***, *t*) and radial growth multiplier *ϑ*^h^. The remodeled geometry and fibrosis distribution are transferred to Model 2, which couples a FitzHugh-Nagumo monodomain model of electrical slow waves *ϕ* with active smooth-muscle tension *T*_act_ and mechanoelectrical feedback. The design of experiments considers five disease-associated factors: hypertrophic remodeling (*ϑ*^h^), which determines the geometry generated by Model 1, and four Model 2 electromechanical factors: electrical diffusivity (*η*_*D*_), excitation threshold (*η*_*a*_), SMC contractility (*η*_*T*_), and wall shear modulus (*η*_*µ*_). Main effects and two-factor interactions are quantified using the motility metric MM.

### 2.2 Inflammation-driven growth model

#### 2.2.1 Fibrosis field evolution

Crohn’s fibrosis originates at focal inflammatory foci and spreads outward through diffusive TGF-*β*/paracrine signaling and myofibroblast activation, saturating once the tissue is fully remodeled (Rieder and Fiocchi 2009; D’Alessio et al. 2022). This focal-source, self-limiting front dynamic is the canonical biological setting for a Fisher-KPP logistic reaction-diffusion equation (Fisher 1937; Sherratt and Murray 1990); we therefore describe the fibrosis state as a dimensionless scalar field *m* ∈ [0, 1], where *m* = 0 denotes fully healthy tissue and *m* = 1 denotes fully fibrotic tissue, whose evolution is governed by

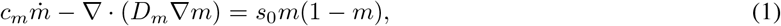

where (.) denotes the material time derivative. Unless otherwise stated, all field gradients and divergences are evaluated in the current configuration, i.e., ***x*** and () ***x*** (). Here *c*_*m*_ is the temporal scaling parameter (analogous to heat capacity), *D*_*m*_ is the isotropic diffusivity, and *s*_0_ is the source rate, with time expressed in decades to represent the chronic timescale of fibrotic remodeling. The logistic right-hand side drives *m* monotonically toward unity and vanishes identically when *m* = 1, capturing the self-limiting saturation of fibrotic remodeling.

Because Abaqus solves the fibrosis PDE by analogy with the transient heat equation, the terms governing fibrosis accumulation, spatial spread, and local progression are implemented using the corresponding thermal quantities (Table 1). The internal energy-like term *U* represents the local accumulation of the fibrosis field, while the diffusive flux ***q*** represents its spatial spread through the tissue. These quantities, together with their consistent derivatives, are returned through UMATHT. The source term S represents the local progression of fibrosis and is returned through UMAT as the heat-source equivalent RPL, together with its linearization DRPLDT and DRPLDE (Dassault Systèmes 2024). Thus, although Abaqus uses thermal terminology, the storage, flux, and source quantities correspond in the present model to local fibrosis accumulation, spatial spread, and local progression, respectively. The storage group encodes the temporal accumulation of the fibrosis field,

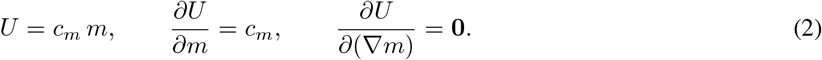

The flux group encodes Fickian diffusion of the fibrosis scalar,

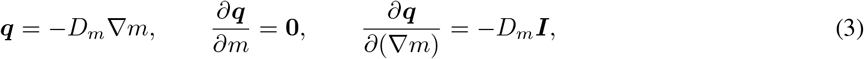

where ***I*** is the second-order identity tensor. The source group encodes the logistic reaction term,

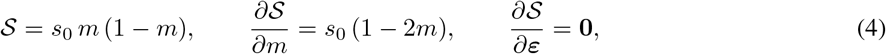

where *S* is the volumetric source rate (returned in RPL) and *∂S/∂m* is its consistent linearization (returned in DRPLDT), required by Abaqus to form the Newton-Raphson tangent for the scalar-field equation. The strain sensitivity *∂S/∂****ε*** = **0** reflects that the fibrosis reaction rate depends only on the scalar field *m* and not on the mechanical deformation state.

#### 2.2.2 Growth law

Let Ω_0_ ⊂ ℝ ^3^ denote the reference configuration, with material coordinates ***X*** ∈ Ω_0_. The motion of the body is described by the deformation map

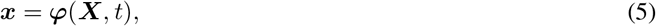

where ***x*** denotes the current spatial position of the material point ***X*** at time *t*. The deformation gradient and the associated kinematic measures are defined as

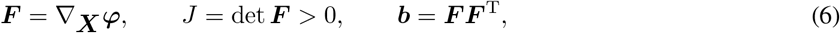

where ***F*** is the deformation gradient, *J* is its determinant (the volumetric Jacobian), and ***b*** is the left Cauchy-Green deformation tensor. Let {***N*** _r_, ***N*** _c_, ***N*** _*ℓ*_}denote a fixed orthonormal basis aligned with the radial, circumferential, and longitudinal directions of the reference configuration, respectively. To account for the irreversible volumetric changes accompanying inflammation-driven wall remodeling within a finite-deformation setting, we adopt the multiplicative decomposition of the deformation gradient into elastic and growth (here, hypertrophy) parts (Rodriguez et al. 1994; Menzel and Kuhl 2012),

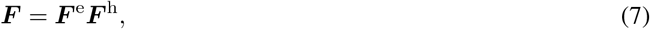

where ***F*** ^e^ is the elastic part and ***F*** ^h^ is the hypertrophic growth part, encoding irreversible structural remodeling. Fibrostenotic Crohn’s strictures are histologically characterized by transmural smooth-muscle hyperplasia and hypertrophy producing predominantly concentric wall thickening with negligible longitudinal elongation (Chen et al. 2017; Koh et al. 2001). This motivates the specific choice of growth tensor,

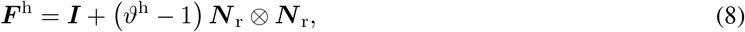

where *ϑ*^h^ is the radial growth multiplier and ***N*** _r_ ⊗ ***N*** _r_ is the structural tensor aligned with the radial direction. The multiplier is driven by the time-varying fibrosis field *m* through the first-order ODE

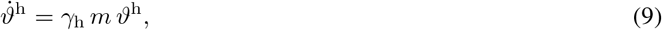

with growth rate *γ*_h_ *>* 0, integrated exactly over each time increment as *ϑ*^h^(*t* + Δ*t*) = *ϑ*^h^(*t*) exp(*γ*_h_*m* Δ*t*). The grown geometry and fibrosis distribution *m* at the end of Model 1 define the reference configuration Ω_0_ for Model 2.

Because Model 1 is used only to generate a representative remodeled geometry, the temporal scaling parameter *c*_*m*_, and the reaction rate *s*_0_ are each set to unity for simplicity (Table 2). The growth rate parameter *γ*_h_ is then calibrated so that the simulation yields a final radial growth multiplier of approximately *ϑ*^h^ = 2.0 in regions of fully developed fibrosis (*m* = 1) after one decade, consistent with the roughly twofold muscular thickening reported in Crohn’s strictures (Chen et al. 2017).

**Table 2.** Parameters for the inflammation-driven growth model (Model 1), governing the reaction-diffusion evolution of the fibrosis field m (Eq. 1) and the inflammation-driven growth ODE (Eq. 9).

| Symbol | Description | Value | Units |
| --- | --- | --- | --- |
| $c_m$ | Temporal scaling parameter (analogous to heat capacity) | 1.0 | — |
| $D_m$ | Fibrosis field diffusivity | 20.0 | mm<sup>2</sup>/decade |
| $s_0$ | Reaction rate parameter | 1.0 | 1/decade |
| $\gamma_h$ | Growth rate parameter | 0.69 | 1/decade |

### 2.3 Mechanical constitutive response of isotropic neo-Hookean matrix with anisotropic fiber reinforcement

The intestinal wall is histologically organized as two layers of smooth muscle, an inner circular layer and an outer longitudinal layer, reinforced by a cross-ply submucosal collagen network (Gabella 1987), with a markedly anisotropic passive response documented under uniaxial and biaxial loading of human and animal gut (Egorov et al. 2002; Bellini et al. 2011). We therefore reuse the constitutive model introduced in our previous study (Anantha Krishnan et al. 2026b): a compressible neo-Hookean ground matrix reinforced by two tension-only fiber families aligned with the circumferential and longitudinal directions (Ciarletta et al. 2009; Carniel et al. 2014)

#### 2.3.1 Cauchy stress

The isotropic strain-energy density is

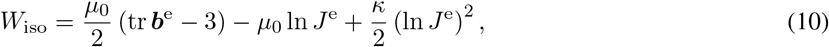

where *µ*_0_ is the infinitesimal shear modulus, *J*^e^ = det ***F*** ^e^ is the elastic Jacobian, ***b***^e^ = ***F*** ^e^(***F*** ^e^)^T^ is the elastic left Cauchy-Green tensor, and *κ* = *κ*_fac_*µ*_0_ is the bulk modulus; setting *κ*_fac_ = 100 enforces a near-incompressibility constraint. The isotropic (ground-matrix) part of the Cauchy stress is

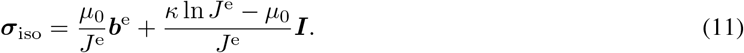

The current unit fiber directions are obtained by pushing the reference directions forward through ***F*** ^e^ and normalizing,

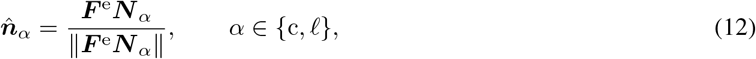

where ***N*** _*α*_ is the reference fiber direction, and the subscripts c and *ℓ* denote the circumferential and longitudinal fiber families, respectively. The corresponding elastic fiber invariants, measuring the squared elastic stretch of each family, are defined as

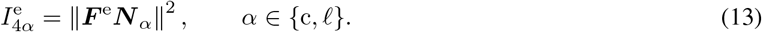

Smooth-muscle fibers carry tension but buckle under compression; we therefore adopt a tension-only polynomial strain-energy density for each fiber family,

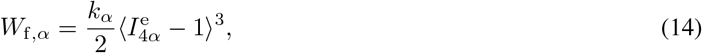

where ⟨·⟩ = max(, 0) enforces tension-only response and *k*_*α*_ is the fiber stiffness of family *α*. The corresponding Cauchy stress contribution of fiber family *α* is

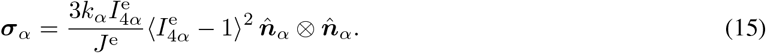

Summing the isotropic ground-matrix contribution and the two fiber families, the total Cauchy stress is

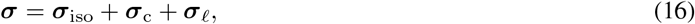

where ***σ***_c_ and ***σ***_*ℓ*_ are the Cauchy stress contributions from the circumferential and longitudinal fiber families, respectively.

#### 2.3.2 Consistent spatial tangent

To form the Newton-Raphson tangent required by Abaqus, the consistent spatial tangent modulus **c** is decomposed into four additive contributions,

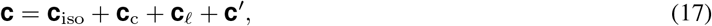

where **c**_iso_ is the isotropic tangent contribution, **c**_c_ and **c**_*ℓ*_ are the circumferential and longitudinal fiber tangent contributions, respectively, and **c***′* is the geometric (initial-stress) tangent. The isotropic contribution takes the form

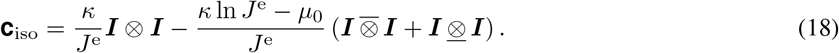

The fiber tangents for the circumferential (*c*) and longitudinal (*ℓ*) families are

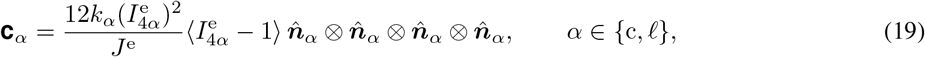

where *k*_c_ and *k*_*ℓ*_ are the stiffness parameters of the circumferential and longitudinal fiber families, respectively, 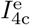 and 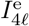 are their corresponding elastic fiber invariants, and 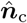 and 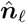 are their current unit directions. Finally, the geometric tangent is given by

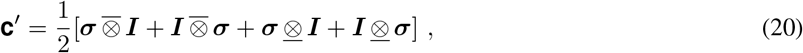

where the fourth-order tensor products are defined componentwise as

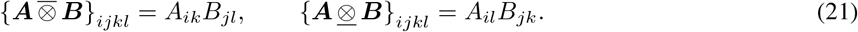

Because *m* influences the Cauchy stress through the growth multiplier *ϑ*^h^, the sensitivity *∂****σ****/∂m* is required for the Newton-Raphson tangent. The growth multiplier is updated through the discrete growth law, so the resulting stress sensitivity corresponds to the algorithmic derivative of the discrete update. Applying the chain rule therefore introduces the factor *γ*_h_Δ*t*. The sensitivity of the isotropic Cauchy stress is

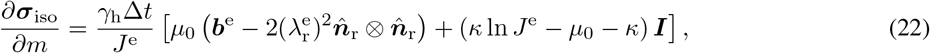

where

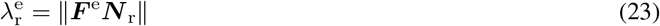

is the elastic radial stretch and 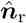 is the current radial unit direction, obtained as in Eq. (12). For each fiber family, the sensitivity takes the form

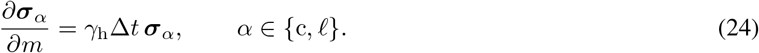

### 2.4 Geometry, mesh, and boundary conditions

The intestinal wall (terminal ileum) is idealized as a quarter-cylinder, exploiting two planes of symmetry in the circumferential direction (Figure 3A and B). Because the geometry, material architecture, fibrotic region, and boundary conditions are symmetric about these planes, the quarter-domain reduces computational cost while retaining the corresponding three-dimensional symmetric response. The geometry is characterized by an inner radius of 8.5 mm, a wall thickness of 1.0 mm. We simulate a short ileal segment (Pierro et al. 2023) with an axial length of 72 mm, four times the diameter, in order to accommodate at least one complete contractile wave. The mesh consists of C3D8T elements with a nominal edge length of 0.5 mm, selected based on convergence of the slow-wave propagation velocity (Brandstaeter et al. 2018). The inflamed region is assumed to occupy the middle 30 % of the axial length and is initialized with *m* = 1 as an initial condition; the remainder of the tube is initialized with *m* = 0 (Figure 3A).

**Figure 3.**
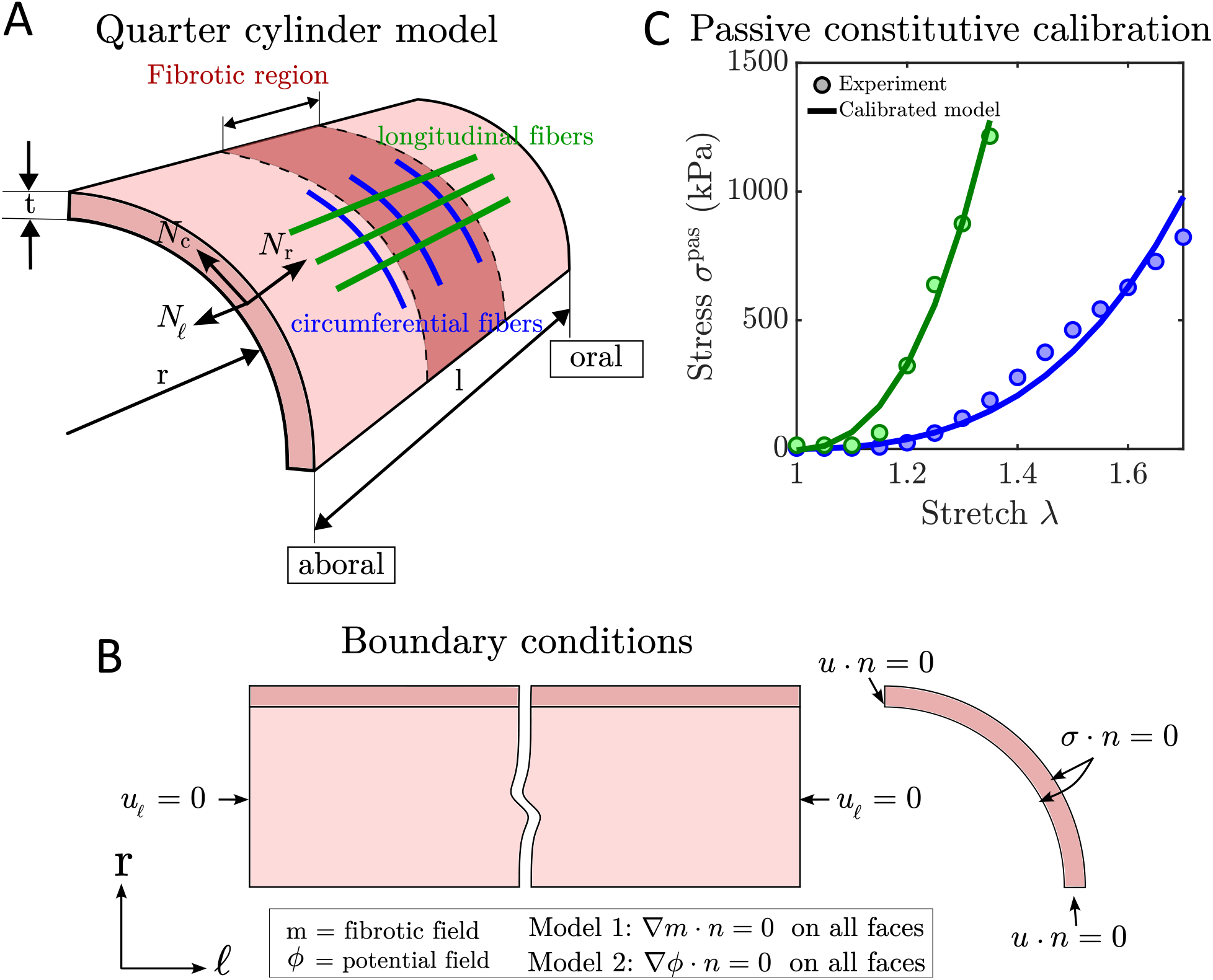
Geometry, boundary conditions, and passive constitutive calibration for the computational framework. **(A)** Quarter-cylinder model of the ileal wall segment, showing the fibrotic region (red), local material basis *{****N*** _r_, ***N*** _c_, ***N*** _*ℓ*_*}* and the embedded circumferential (blue) and longitudinal (green) fiber families. The quarter-cylinder exploits two symmetry planes to reduce computational cost while preserving the full three-dimensional stress state. **(B)** Boundary conditions applied to both models. Axial displacement is constrained on the proximal and distal faces (*u*_*ℓ*_ = 0). The two cut faces enforce symmetry (***u · n*** = 0), and the inner lumen and outer surfaces are traction-free (***σ****·****n*** = 0). Zero flux (*∇ m·****n*** = 0 and *∇ ϕ·****n*** = 0) is the natural boundary condition for the scalar fields in both models. **(C)** Passive constitutive calibration of the fiber-reinforced model against uniaxial extension data from Egorov et al. (2002) for the circumferential (blue) and longitudinal (green) smooth muscle directions. The calibrated fiber stiffnesses *k*_c_ and *k*_*ℓ*_ reproduce the measured nonlinear stiffening in both directions simultaneously.

Symmetry boundary conditions are applied on the two cut faces of the quarter-cylinder (***u****·****n*** = 0, where ***n*** is the outward unit normal). The proximal and distal faces are constrained axially (*u*_*ℓ*_ = 0, where *u*_*ℓ*_ is the displacement component in the longitudinal direction) but free in the radial and circumferential directions. The inner and outer surfaces are traction-free (***σ · n*** = **0**) in both models. In Model 1, zero fibrosis flux (∇ *m* · ***n*** = 0) holds on all faces, satisfied naturally by the UMATHT formulation. In Model 2, the zero electrical flux (∇ *ϕ* · ***n*** = 0) holds on all faces for all time, consistent with the natural boundary condition of the monodomain (a single conductive domain, without separately resolving the intracellular and extracellular spaces) formulation (Figure 3B).

The parameters *k*_c_ and *k*_*ℓ*_ were tuned against the experimental curves, with root mean square error used to assess the fit (Figure 3C).

### 2.5 Electromechanical model

Model 2 describes the electromechanical behavior on either the non-hypertrophied or hypertrophically remodeled reference geometry, according to the DOE level of *ϑ*^h^. The normalized ICC potential field *ϕ* is solved as the primary scalar field via a UMATHT, while a UMAT controls passive mechanics, active stress generation, and mechanoelectrical feedback. The fibrosis field *m* is imported as a predefined field. For the hypertrophied DOE level, the grown geometry obtained from Model 1 serves as the reference configuration for Model 2; for the non-hypertrophied level, the corresponding geometry with *ϑ*^h^ = 1 is used. No further growth occurs during the electromechanical simulation, so ***F*** ^h^ = ***I*** within Model 2.

In the inflamed and fibrotic tissue, four dimensionless scaling parameters (*η*_*D*_, *η*_*a*_, *η*_*T*_, *η*_*µ*_) alter the electrical, contractile, and passive mechanical response to represent the principal non-geometric effects of disease. Of the two electrical factors, *η*_*D*_ reduces the effective electrical diffusivity and thereby alters slow-wave conduction, whereas *η*_*a*_ lowers the FitzHugh-Nagumo excitation threshold and modifies local excitability. Both alterations are scaled spatially by the fibrosis field *m*(***x***) rather than by imposing a prescribed conduction block. The four scaling parameters are active exclusively in Model 2, so the remodeled geometry established by Model 1 is independent of them. Each factor is introduced with its governing relation below, and baseline values for all electromechanical parameters are listed in Table 3.

**Table 3.**
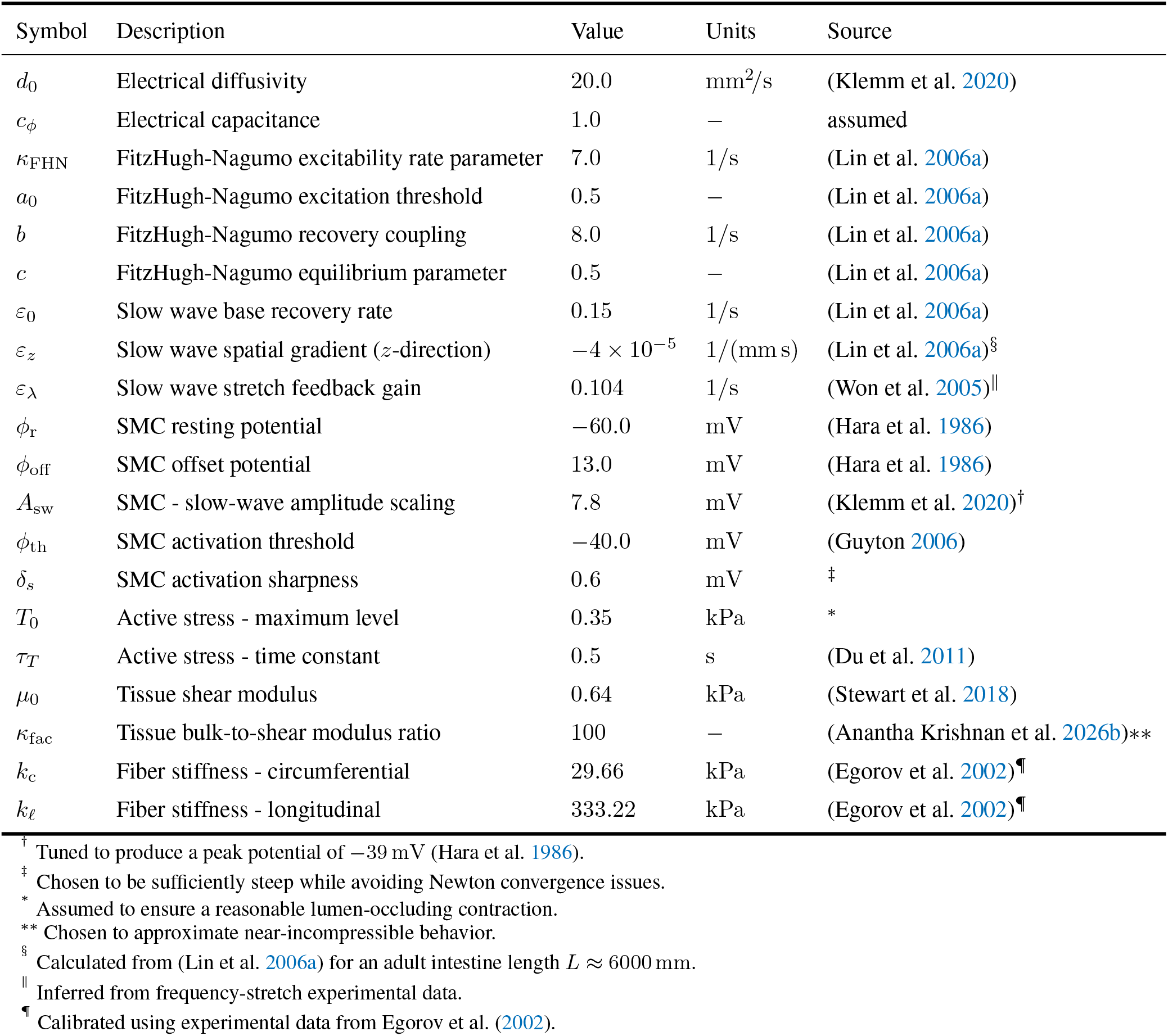
Electromechanical parameters for Model 2. Parameters governing FitzHugh-Nagumo kinetics, SMC electrophysiology, and passive mechanics are listed with their physiological sources.

| Symbol | Description | Value | Units | Source |
| --- | --- | --- | --- | --- |
| $d_0$ | Electrical diffusivity | 20.0 | $\text{mm}^2/\text{s}$ | (Klemm et al. 2020) |
| $c_\phi$ | Electrical capacitance | 1.0 | — | assumed |
| $\kappa_{\text{FHN}}$ | FitzHugh-Nagumo excitability rate parameter | 7.0 | 1/s | (Lin et al. 2006a) |
| $a_0$ | FitzHugh-Nagumo excitation threshold | 0.5 | — | (Lin et al. 2006a) |
| $b$ | FitzHugh-Nagumo recovery coupling | 8.0 | 1/s | (Lin et al. 2006a) |
| $c$ | FitzHugh-Nagumo equilibrium parameter | 0.5 | — | (Lin et al. 2006a) |
| $\varepsilon_0$ | Slow wave base recovery rate | 0.15 | 1/s | (Lin et al. 2006a) |
| $\varepsilon_z$ | Slow wave spatial gradient ($z$-direction) | $-4 \times 10^{-5}$ | $1/(\text{mm s})$ | (Lin et al. 2006a)<sup>§</sup> |
| $\varepsilon_\lambda$ | Slow wave stretch feedback gain | 0.104 | 1/s | (Won et al. 2005) |
| $\phi_r$ | SMC resting potential | -60.0 | mV | (Hara et al. 1986) |
| $\phi_{\text{off}}$ | SMC offset potential | 13.0 | mV | (Hara et al. 1986) |
| $A_{\text{sw}}$ | SMC - slow-wave amplitude scaling | 7.8 | mV | (Klemm et al. 2020)<sup>†</sup> |
| $\phi_{\text{th}}$ | SMC activation threshold | -40.0 | mV | (Guyton 2006) |
| $\delta_s$ | SMC activation sharpness | 0.6 | mV | <sup>‡</sup> |
| $T_0$ | Active stress - maximum level | 0.35 | kPa | * |
| $\tau_T$ | Active stress - time constant | 0.5 | s | (Du et al. 2011) |
| $\mu_0$ | Tissue shear modulus | 0.64 | kPa | (Stewart et al. 2018) |
| $\kappa_{\text{fac}}$ | Tissue bulk-to-shear modulus ratio | 100 | — | (Anantha Krishnan et al. 2026b)** |
| $k_c$ | Fiber stiffness - circumferential | 29.66 | kPa | (Egorov et al. 2002)<sup>¶</sup> |
| $k_\ell$ | Fiber stiffness - longitudinal | 333.22 | kPa | (Egorov et al. 2002)<sup>¶</sup> |

<sup>†</sup> Tuned to produce a peak potential of -39 mV (Hara et al. 1986).
<sup>‡</sup> Chosen to be sufficiently steep while avoiding Newton convergence issues.
\*
Assumed to ensure a reasonable lumen-occluding contraction.
\*\* Chosen to approximate near-incompressible behavior.
<sup>§</sup> Calculated from (Lin et al. 2006a) for an adult intestine length $L \approx 6000$ mm.
<sup>||</sup> Inferred from frequency-stretch experimental data.
<sup>¶</sup> Calibrated using experimental data from Egorov et al. (2002).

#### 2.5.1 Governing PDE

ICC form a gap-junction-coupled excitable syncytium dominated by intracellular conduction. Consequently, tissue-scale electrical activity is well-described by a monodomain reaction-diffusion formulation (Du et al. 2018). We adopt the FitzHugh-Nagumo reaction term in place of more biophysically detailed ionic models because it effectively reproduces the key features of intestinal slow-wave dynamics, including threshold excitability, refractoriness, conduction velocity, and frequency-gradient entrainment, while remaining computationally tractable for three-dimensional finite-element electromechanical simulations (Aliev et al. 2000; Lin et al. 2006a; Klemm et al. 2020). The normalized ICC potential *ϕ* ∈ [0, 1] satisfies

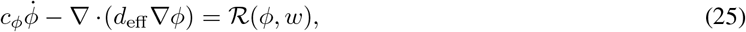

where *c*_*ϕ*_ is the electrical capacitance, *d*_eff_ is the effective electrical diffusivity, *w* is the FitzHugh-Nagumo recovery variable, and *R* is the ionic source term. ICC depletion, driven primarily by the inflammatory microenvironment with progressive fibrosis acting as an additional compounding factor, reduces the local conduction velocity via the effective diffusivity

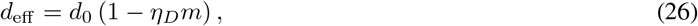

where *d*_0_ is the healthy baseline diffusivity and *η*_*D*_ ∈ [0, 1] is the dimensionless diffusivity scaling parameter. The monodomain relation 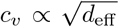 inherited from continuous-cable theory (Kléber and Rudy 2004) means that *η*_*D*_ can be calibrated directly against measured conduction-velocity changes, linking this tissue-level parameter to an experimentally accessible quantity (Table 4). Following the same UMATHT structure as Model 1 (Table 1), the storage group encodes the temporal accumulation of the electrical potential,

**Table 4.**
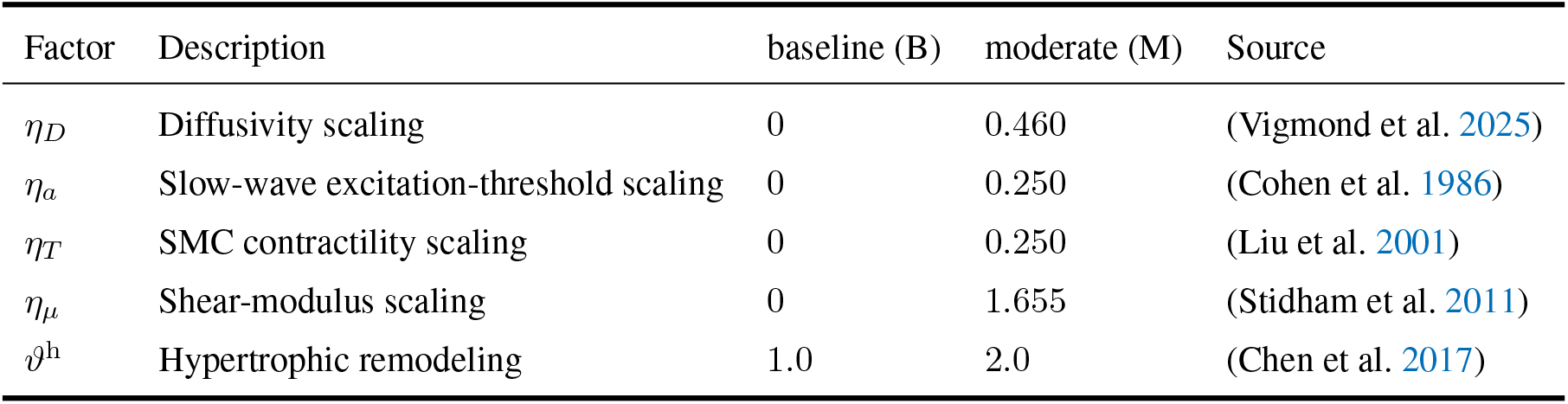
Factors and levels used in the full-factorial 2^5^ design of experiments. For the four Model 2 electromechanical scaling parameters, the baseline (B) level corresponds to *η* = 0 and the moderate-disease (M) level to *η* = 0.5 *η*_max_. For Model 1 hypertrophic remodeling, B and M correspond to *ϑ*^h^ = 1.0 and 2.0, respectively.

| Factor | Description | baseline (B) | moderate (M) | Source |
| --- | --- | --- | --- | --- |
| $\eta_D$ | Diffusivity scaling | 0 | 0.460 | (Vigmond et al. 2025) |
| $\eta_a$ | Slow-wave excitation-threshold scaling | 0 | 0.250 | (Cohen et al. 1986) |
| $\eta_T$ | SMC contractility scaling | 0 | 0.250 | (Liu et al. 2001) |
| $\eta_\mu$ | Shear-modulus scaling | 0 | 1.655 | (Stidham et al. 2011) |
| $\vartheta^h$ | Hypertrophic remodeling | 1.0 | 2.0 | (Chen et al. 2017) |

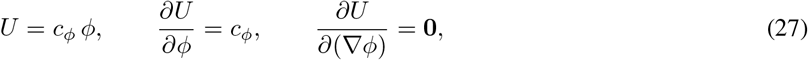

where *U* is the internal energy-like storage term and *c*_*ϕ*_ is the electrical capacitance (analogous to specific heat capacity in the Abaqus heat-transfer analogy). The flux group encodes Fickian conduction of the normalized potential,

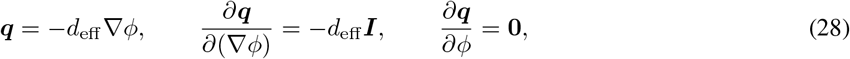

where the zero term (DFDT) reflects that the flux has no direct dependence on the ICC potential *ϕ*, because the effective diffusivity *d*_eff_ depends only on the predefined fibrosis field *m*.

#### 2.5.2 FitzHugh-Nagumo ionic kinetics

The ionic source term follows the FitzHugh-Nagumo formulation (Aliev et al. 2000; Lin et al. 2006a),

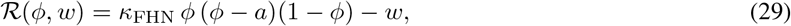

where *a* is the excitation threshold, *κ*_FHN_ is the excitability rate parameter, and *w* is a slow recovery variable satisfying the ODE

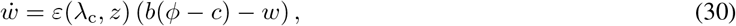

where *ε* is the recovery rate, driven by mechanoelectrical feedback; *λ*_c_ is the circumferential stretch; *z* is the longitudinal coordinate; *b* is the FitzHugh-Nagumo recovery coupling parameter; and *c* is the FitzHugh-Nagumo equilibrium parameter. Inflammation depolarizes the smooth-muscle resting membrane potential, reducing the excitation gap and altering tissue excitability (Cohen et al. 1986); we represent this by a linear reduction of the excitation threshold,

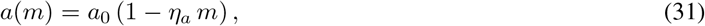

where *a*_0_ is the healthy baseline excitation threshold and *η*_*a*_ is the dimensionless threshold scaling parameter.

#### 2.5.3 Ionic source term linearization

The ionic source term is linearized consistently with the backward-Euler time integration of the recovery variable. The resulting algorithmic Jacobian is

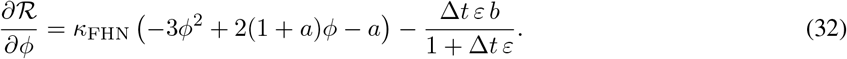

This quantity is returned through the Abaqus `DRPLDT` and is used to assemble the electrical contribution to the coupled Newton-Raphson tangent.

#### 2.5.4 Mechanoelectrical feedback

ICC and intestinal smooth muscle express stretch-activated ion channels whose effect on slow-wave frequency and refractoriness has been demonstrated directly in gastric muscle (Won et al. 2005) and documented in the gastrointestinal tract broadly (Kraichely and Farrugia 2007). The recovery rate is therefore assumed to depend on local stretch, such that increased circumferential stretch accelerates recovery and thereby shortens the action-potential duration. We let *ε* be a function of the circumferential stretch *λ*_c_ and the current spatial coordinate *z* along the longitudinal axis of the intestinal segment,

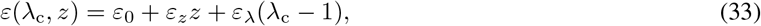

where *ε*_0_ is the spatially uniform baseline recovery rate, *ε*_*z*_ is the longitudinal gradient of the recovery rate, and *ε*_*λ*_ is the stretch feedback gain. To prevent division by zero in the Newton-Raphson tangent, *ε* is floored at 10^*−*12^ in the implementation. This form embeds mechanoelectrical feedback into the reaction term in line with established practice in gastric electromechanics (Klemm et al. 2020).

### 2.6 Active stress model

#### 2.6.1 Smooth muscle cell potential

The potential of the SMC is obtained from the dimensionless field *ϕ* via the linear mapping

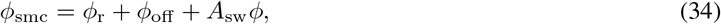

where *ϕ*_smc_ is the dimensional membrane potential of the SMC, *ϕ*_r_ is the SMC resting membrane potential, *ϕ*_off_ is a constant offset, and *A*_sw_ is the electrochemical coupling gain due to slow waves.

#### 2.6.2 Activation gate

The steady-state force-[Ca^2+^] relationship in smooth muscle is a cooperative, sigmoidal curve reflecting cross-bridge recruitment (Rembold and Murphy 1990; DeFeo and Morgan 1985). We therefore model the dimensionless activation gate as a logistic function of the SMC membrane potential,

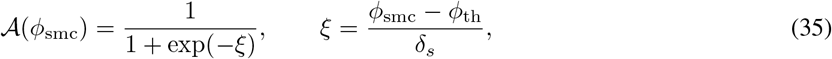

where *A* is the dimensionless activation gate (ranging from 0 to 1), *ξ* is the dimensionless argument of the logistic function, *ϕ*_th_ is the half-activation threshold and *δ*_*s*_ controls the steepness of the transition, consistent with Hill-type activation kernels adopted in gastrointestinal smooth-muscle continuum models (Gajendiran and Buist 2011; Klemm et al. 2020).

#### 2.6.3 Active tension evolution

Inflammation reduces L-type calcium channel function and expression, thereby decreasing peak active tension (Liu et al. 2001; Shi and Sarna 2005),

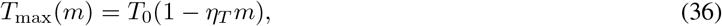

where *T*_0_ is the healthy baseline peak active stress and *η*_*T*_ is the dimensionless contractility scaling parameter. Because the chemo-mechanical cascade linking SMC depolarization to cross-bridge force introduces a finite delay between electrical activation and force generation, as formalized by Hai-Murphy latch-bridge kinetics (Hai and Murphy 1988; Murtada et al. 2010), we model the active tension by a first-order ODE,

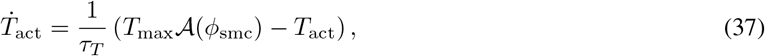

where *T*_act_ is the current active tension and *τ*_*T*_ is the active stress time constant. This form is in line with phenomenological active-stress formulations established for visceral and vascular smooth muscle (Rachev and Hayashi 1999; Stålhand et al. 2008) and with gastrointestinal electromechanical models that adopt analogous activation kinetics (Du et al. 2011; Klemm et al. 2020).

#### 2.6.4 Active Cauchy stress

Intestinal circular smooth muscle shortens primarily in the circumferential direction during propagating contraction (Gabella 1987; Egorov et al. 2002); accordingly, active stress is assumed to act exclusively along the current circumferential direction 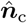

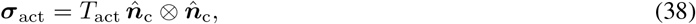

where ***σ***_act_ is the active Cauchy stress tensor and 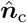 is the current unit circumferential direction. The consistent stress-potential coupling tangent returned through DDSDDT is

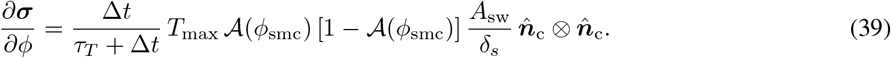

### 2.7 Passive constitutive law and coupled tangent

#### 2.7.1 Passive mechanics

The passive constitutive law in Model 2 is identical in form to that of the growth model (Sections 2.2-2.3.2), with two modifications. First, there is no growth (***F*** ^h^ = ***I***, hence ***F*** ^e^ = ***F*** and *J*^e^ = *J* = det ***F***, and all invariants are calculated using the total deformation gradient). Second, the shear modulus is increased to represent the altered mechanical properties arising from structural remodeling in Crohn’s strictures, consistent with elastographic measurements (Stidham et al. 2011; Stewart et al. 2018),

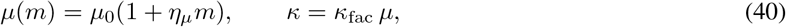

where *µ*(*m*) is the fibrosis-dependent shear modulus and *η*_*µ*_ is the dimensionless shear-modulus scaling parameter. The passive Cauchy stress follows directly from the neo-Hookean and fiber expressions of Section 2.3 evaluated at ***F*** ^e^ = ***F***. The total Cauchy stress in Model 2 is the sum of the passive and active contributions,

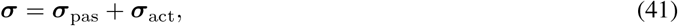

where ***σ***_pas_ = ***σ***_iso_ + ***σ***_c_ + ***σ***_*ℓ*_.

#### 2.7.2 Consistent spatial tangent

The total spatial tangent for Model 2 is assembled as the sum of passive, active, and geometric contributions,

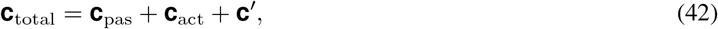

where **c**_pas_ is the passive tangent consisting of isotropic, circumferential, and longitudinal fiber contributions, **c**_act_ is the active tangent contribution, and **c***′* is the geometric tangent. The passive contribution **c**_pas_ follows the same form as Section 2.3.2 with *µ*(*m*) replacing *µ*_0_ and *J* replacing *J*^e^. Letting 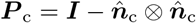 denote the projection onto the plane normal to the circumferential direction 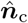 the active tangent contribution is

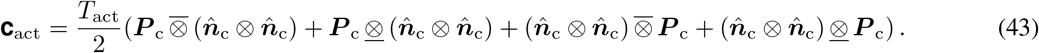

#### 2.7.3 Source-term sensitivity to strain

Mechanoelectrical feedback introduces an implicit dependence of the ionic source term *R* on the circumferential stretch *λ*_c_ through the recovery rate *ε*(*λ*_c_). This is given by

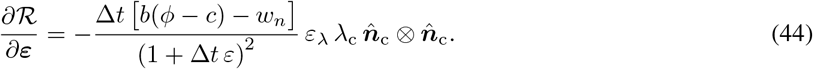

This quantity is returned to Abaqus through DRPLDE and provides the strain sensitivity of the ionic source term.

### 2.8 Model coupling and data transfer

Upon completion of Model 1, the final time step of the simulation is post-processed using a Python script, which reads the deformed nodal coordinates and exports ten predefined field variables: the fibrosis scalar *m*, and the three components each of the local radial, circumferential, and longitudinal unit vectors. These unit vectors are recomputed geometrically from the deformed nodal positions used in Model 1, ensuring the triad reflects the grown geometry transferred to Model 2. These fields are written as an Abaqus *INITIAL CONDITIONS, TYPE=FIELD include file and are read directly by Model 2 at the start of its analysis. The grown mesh from Model 1 serves as the reference configuration for Model 2, with ***F*** ^h^ = ***I*** enforced throughout. It is important to note that only the geometry and field variables are transferred. The residual stresses developed during growth are not imported, and the remodeled configuration is adopted as a stress-free reference for Model 2, consistent with the assumption that remodeling occurs slowly enough for the tissue to relax between the two stages.

### 2.9 Motility metric

propagating contraction produces cyclic changes in lumen volume. Clinically, quantitative cine-MRI approaches assess intestinal motility from the temporal variation in bowel deformation; for example, Menys et al. (2013a) quantify local expansion and contraction using the standard deviation of the Jacobian determinant obtained from non-rigid registration of sequential MR images. Following the same general principle of using temporal deformation variability as a surrogate for motility, we define the motility metric as the standard deviation of lumen volume over one complete contraction cycle,

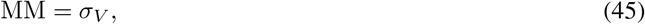

where *σ*_*V*_ is the population standard deviation of the lumen-volume signal *V* (*t*) over the selected cycle. Accordingly, MM has units of volume mm^3^ and quantifies the absolute magnitude of cyclic lumen-volume variation. A larger MM indicates greater cyclic lumen deformation, whereas a smaller value indicates reduced cyclic deformation. The lumen volume is computed post hoc from the Model 2 output database at each output frame. Cross-sectional areas of the lumen are calculated from angularly ordered inner-surface nodes using the shoelace formula and then integrated along the axial direction using the trapezoidal rule. The same axial region is evaluated in all simulations. Because *V* (*t*) is integrated over the same axial region in every simulation, MM represents segment-level cyclic lumen deformation rather than local occlusion at a single cross-section. The metric does not quantify transport of luminal contents or propulsive efficiency. The dominant contraction period is estimated from the lumen-volume signal using the fast Fourier transform, and the final complete contraction cycle is identified from successive volume minima separated by approximately this period. The selected cycle is then interpolated onto 1000 equally spaced time points, from which *σ*_*V*_ is calculated.

### 2.10 Design of experiments and statistical analysis

Taken together, the five DOE factors represent selected electrical, contractile, mechanical, and structural consequences of fibrostenosing Crohn’s disease. The parameter *η*_*D*_ reduces electrical diffusivity (Eq. 26), *η*_*a*_ lowers the FitzHugh-Nagumo excitation threshold (Eq. 31), *η*_*T*_ reduces peak active tension (Eq. 36), and *η*_*µ*_ increases the wall shear modulus (Eq. 40). The fifth factor, *ϑ*^h^, represents structural hypertrophic remodeling and distinguishes the healthy geometry (*ϑ*^h^ = 1) from the remodeled geometry generated by Model 1 (*ϑ*^h^ ≈ 2 in the fibrotic region). A full-factorial 2^5^ DOE is used to estimate the main effects and all two-factor interactions among the five factors, with each factor independently assigned to its baseline or moderate-disease level. The four electromechanical scaling parameters (*η*_*D*_, *η*_*a*_, *η*_*T*_, *η*_*µ*_) are varied between a baseline level (B, *η* = 0) and a moderate-disease level (M, *η* = 0.5 *η*_max_). Hypertrophic remodeling is varied between *ϑ*^h^ = 1.0 (B; no hypertrophy) and *ϑ*^h^ = 2.0 (M; remodeled geometry). This results in 32 electromechanical simulations. The fibrosis distribution used by Model 2 was held fixed across the two hypertrophic-remodeling levels so that *ϑ*^h^ could be varied independently of the four electromechanical factors. Runs 1–16 use the non-hypertrophied geometry (*ϑ*^h^ = 1), whereas Runs 17–32 use the hypertrophically remodeled geometry generated by Model 1; the same spatial fibrosis field *m*(***x***) is used to scale the Model 2 disease parameters in both geometries. Run 1 therefore represents the reference configuration, with all electromechanical disease parameters at baseline and *ϑ*^h^ = 1.

Of the five disease-associated factors considered in the DOE, four could be parameterized from reported intestinal or Crohn’s disease measurements: excitation threshold, SMC contractility, wall stiffness, and hypertrophic remodeling. For the excitation threshold, the resting membrane potential of healthy SMC *ϕ*_r_ = − 60 mV lies 20 mV below the firing threshold *ϕ*_th_ = −40 mV. Inflammation depolarizes the resting potential by approximately 10 mV (Cohen et al. 1986), halving this excitation gap, which in the dimensionless FitzHugh-Nagumo system corresponds to *a*(*m* = 1) = *a*_0_*/*2 = 0.25 and hence *η*_*a*_ = 0.5. For SMC contractility, the reduction in peak active tension is based on reported impairment of L-type calcium-channel function in inflamed intestinal smooth muscle (Liu et al. 2001). For wall stiffness, the shear-modulus scaling was derived from reported elastography measurements of Crohn’s disease bowel tissue (Stidham et al. 2011). For hypertrophic remodeling, histological measurements indicate approximately twofold muscular thickening in Crohn’s strictures (Chen et al. 2017). We therefore set the moderate-disease level to *ϑ*^h^ = 2.0, while *ϑ*^h^ = 1.0 represents the baseline state with no hypertrophic growth.

Direct intestinal measurements of electrical diffusivity or conduction velocity across graded fibrosis are currently unavailable, so the diffusivity scaling was estimated using cardiac data as a surrogate: dense fibrosis reduces conduction velocity by up to 75 % (Vigmond et al. 2025), corresponding to a healthy velocity *c*_*v*,0_ ≈ 70 cm*/*s and a severely fibrotic velocity *c*_*v*,1_ ≈ 20 cm*/*s. The monodomain relation 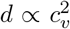 then gives *η*_*D*_ = 1 − (*c*_*v*,1_*/c*_*v*,0_)^2^ ≈ 0.92, providing a practical surrogate for reduced electrical coupling.

Model 1 was executed once for the healthy (baseline) case and once for the fibrotic case. In the healthy case the fibrosis field is zero everywhere, so *ϑ*^h^ = 1 throughout and the geometry is returned unchanged. This run is performed solely so that both cases pass through an identical simulation pipeline, allowing the same post-processing script to export the field variables and the same Model 2 input structure to be used for every simulation. The resulting geometry and fiber fields were then exported and used as inputs for Model 2. Each electromechanical simulation was run for 180 s, after which the slow-wave activity had reached entrainment. Results are reported from the final contraction cycle of each simulation. The motility metric is expressed as the percentage change from the healthy baseline MM_0_,

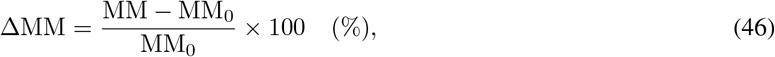

where MM_0_ is the motility metric for Run 1. A linear regression model containing all five main effects and all ten two-factor interactions is fitted to the 32 responses,

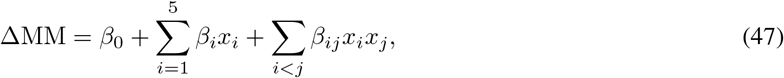

where *x*_*i*_ = −1 and *x*_*i*_ = +1 denote the B and M levels, respectively. Because the factors are coded using {−1, +1}levels, the full effect associated with moving a factor from B to M is 2*β*_*i*_. Regression coefficients are estimated by ordinary least squares and ranked by absolute magnitude.

## 3 Results

### 3.1 Fibrosis field evolution and wall growth

In Model 1 simulations of intestinal growth, the fibrosis scalar field *m* evolved from its initial seed region and propagated outward according to the logistic reaction-diffusion equation, reaching a spatially graded distribution across the fibrotic segment within the simulated time window (Figure 4). The field saturated at *m* ≈ 1 in the core of the fibrotic zone while decaying smoothly to *m* = 0 in the flanking healthy tissue. The radial growth multiplier *ϑ*^h^ increased monotonically with local fibrosis intensity, producing progressive thickening of the fibrotic segment. At maximum fibrosis (*m* = 1), the muscular layer reached approximately twice its undeformed thickness, consistent with the smooth-muscle hypertrophy reported by Chen et al. (2017). The grown geometry, fibrosis distribution, and local fiber triad were exported and transferred to Model 2 as predefined fields.

**Figure 4.**
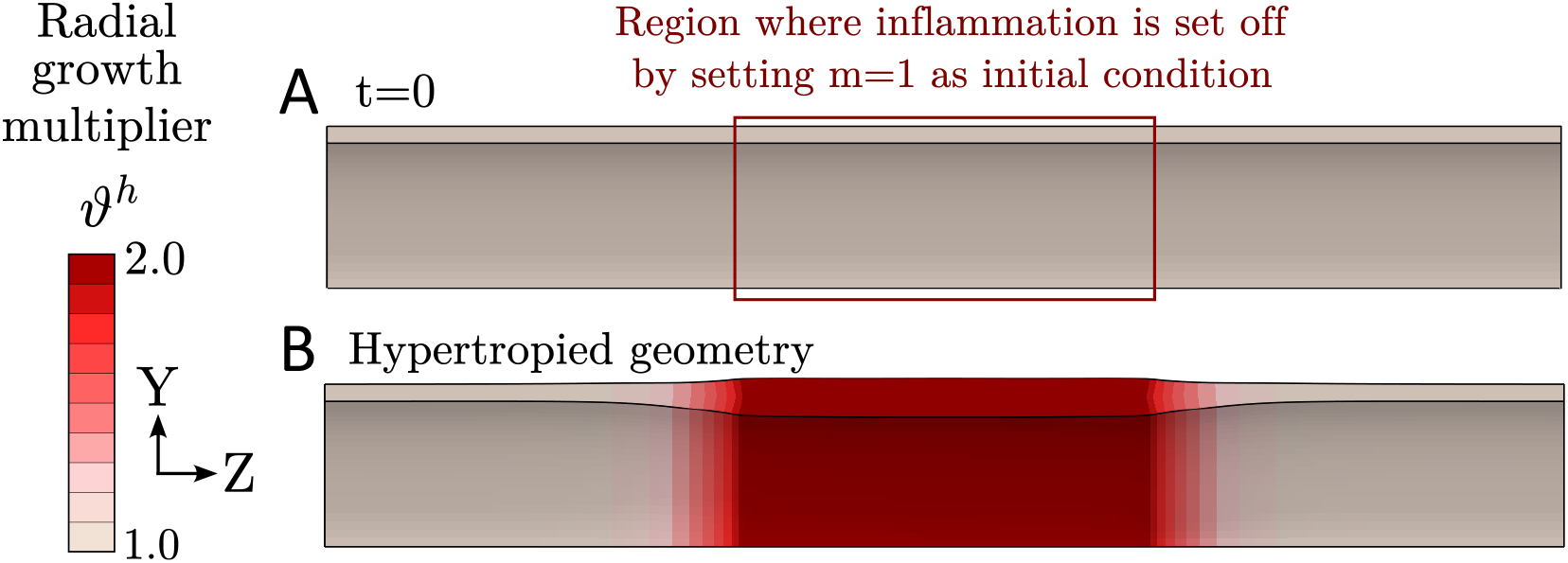
Model 1 results showing inflammation-driven radial wall growth. **(A)** Initial configuration (*t* = 0) with the fibrotic seed region, where *m* = 1 is prescribed as an initial condition over the central 30% of the axial length; the remainder of the domain is initialized to *m* = 0. **(B)** Remodeled geometry at the end of the growth simulation, showing radial wall thickening localized to the fibrotic segment; the growth multiplier reaches a peak value of *ϑ*^h^ = 2.0 at the core of the fibrotic zone, producing a geometry that is transferred as the reference configuration to Model 2. The smooth spatial gradient of *ϑ*^h^ between the fibrotic core and the flanking healthy tissue reflects the diffusive spreading of the fibrosis field *m*.

### 3.2 Healthy baseline electromechanics

In the healthy baseline configuration, corresponding to Run 1 (*η*_*D*_ = *η*_*a*_ = *η*_*T*_ = *η*_*µ*_ = 0 and *ϑ*^h^ = 1), the FitzHugh-Nagumo monodomain model generated a propagating slow wave that traversed the intestinal segment from proximal to distal in approximately 6.6 s. The resulting cyclic lumen-volume signal yielded a baseline motility metric of

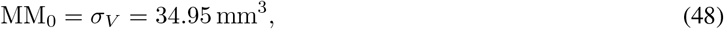

which serves as the reference for all subsequent values of ΔMM. Across the DOE, the cycle duration remained approximately 6.6 s. The smooth-muscle activation gate followed the passage of the wave, producing a circumferential contraction that reasonably occluded the lumen transiently at each axial position (Figure 5). The mechanoelectrical feedback term modulated the recovery rate along the axial direction, producing a frequency gradient consistent with the physiological oro-aboral gradient reported by Lin et al. (2006a).

**Figure 5.**
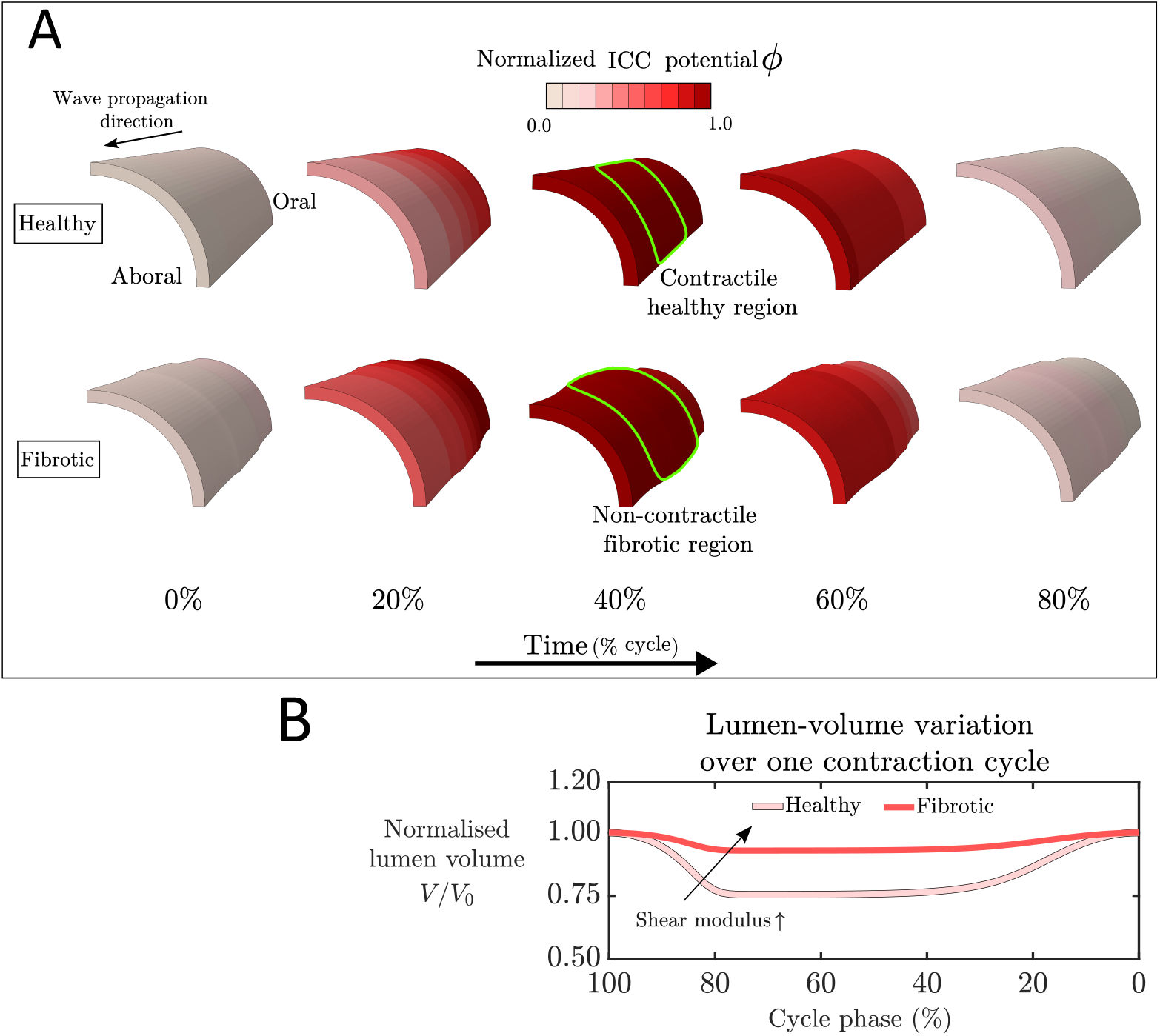
Model 2 electromechanical simulation results for the healthy baseline (Run 1) and the fully perturbed configuration (Run 32), in which all five disease-associated factors are at their moderate-disease level. **(A)** Snapshots of the normalized ICC slow-wave potential *ϕ∈* [0, 1] at five equally spaced instants across one contraction cycle (0, 20, 40, 60 and 80% of cycle phase), for the healthy segment (top) and the fibrotic segment (bottom). A coherent wave of depolarization propagates aborally in both cases, with no conduction block; the cycle duration remained approximately 6.6 s across the DOE. The two rows differ in their mechanical response: at 40 % of cycle phase the healthy segment shortens markedly within the activated region (green outline, top), whereas the fibrotic segment remains close to its reference configuration (green outline, bottom) despite preserved electrical activation. **(B)** Normalized lumen volume *V/V*_0_ over one complete contraction cycle, where *V*_0_ is the volume at cycle onset. The healthy baseline contracts to approximately 75 % of *V*_0_, the fibrotic run only to 93 %.

### 3.3 Effect of hypertrophic remodeling

Hypertrophic remodeling modestly reduced cyclic lumen-volume deformation across the DOE. Averaged over all matched electromechanical parameter combinations, changing the growth state from *ϑ*^h^ = 1 to *ϑ*^h^ = 2 produced a full effect of −1.47 percentage points (pp) in ΔMM, corresponding to an average motility-metric ratio of 0.976. Thus, hypertrophic wall remodeling reduced the absolute magnitude of cyclic lumen-volume deformation by approximately 2.4 % over the parameter range examined. Because MM = *σ*_*V*_ quantifies absolute cyclic lumen-volume deformation, this reduction reflects the combined geometric consequences of hypertrophic wall thickening and luminal narrowing on the magnitude of the cyclic mechanical response.

### 3.4 Fibrotic electromechanics and mechanistic attribution

Across all 32 DOE configurations, slow-wave propagation was preserved through the fibrotic segment. Peak *ϕ* remained detectable throughout the fibrotic region, no conduction block was observed, and the cycle duration remained approximately 6.6 s across the DOE (Figure 5A). Local active stress generation likewise remained present, indicating that motility loss did not arise from loss of electrical activation. The difference between the healthy and fibrotic segments was instead mechanical: at comparable activation, the fibrotic wall underwent markedly less circumferential shortening, and lumen occlusion was correspondingly reduced. Consequently, cyclic lumen-volume variation was markedly attenuated: the healthy baseline decreased to approximately 75 % of the cycle-onset volume, whereas the fibrotic configuration decreased only to approximately 93 % (Figure 5B).

### 3.5 DOE: effect of disease mechanisms on the motility metric

Across the 32 DOE configurations, ΔMM ranged up to −72.03% in Run 15, with a mean response of −40.88% (Figure 6). Every disease-perturbed configuration therefore exhibited a motility metric below the healthy reference over the range examined. Run 15 yielded MM = 9.77 mm^3^, compared with MM_0_ = 34.95 mm^3^ in Run 1. When all five disease factors were simultaneously at their moderate-disease levels (Run 32), the motility metric was 9.92 mm^3^, corresponding to a 71.61 % reduction from baseline.

**Figure 6.**
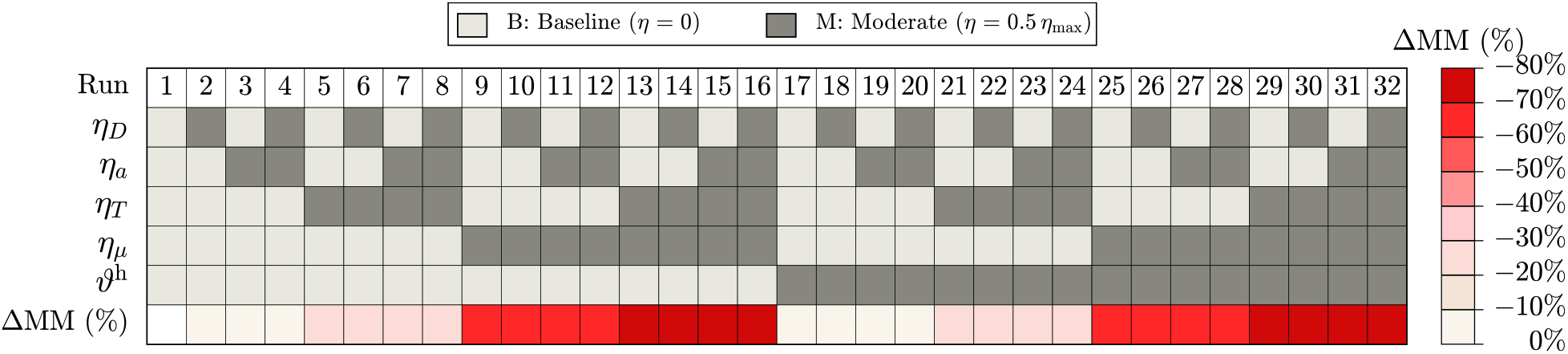
Full-factorial 2^5^ DOE across electrical diffusivity (*η*_*D*_), excitation threshold (*η*_*a*_), peak active stress (*η*_*T*_), wall shear modulus (*ηµ*), and hypertrophic remodeling (*ϑ*^h^). Each column represents one of the 32 experimental runs. Light-grey cells denote the baseline level, whereas dark-grey cells denote the moderate-disease level. For the four electromechanical scaling parameters, baseline corresponds to *η* = 0 and moderate to *η* = 0.5 *η*_max_. For hypertrophy, baseline corresponds to *ϑ*^h^ = 1.0 and moderate to *ϑ*^h^ = 2.0. The bottom row shows the percentage change in motility metric relative to Run 1 (MM_0_ = 34.95 mm^3^); increasingly dark red indicates greater motility loss.

### 3.6 Regression analysis and effect ranking

The regression model containing five main effects and all ten two-factor interactions was fitted to the 32 responses. All five main effects were negative, indicating a reduction in the motility metric when each factor was moved from baseline to moderate-disease level (Figure 7; Figure 8A). Wall shear modulus was the dominant factor, with *β*_*µ*_ = −25.40 pp and a full effect of −50.79 pp. Peak active stress was the second-largest main effect (*β*_*T*_ = −8.28 pp; full effect −16.56 pp). The remaining main effects were substantially smaller: excitation threshold produced a full effect of −1.89 pp, hypertrophic remodeling −1.47 pp, and electrical diffusivity −0.39 pp. Based on the absolute magnitudes of the full main effects, the resulting ranking was *η*_*µ*_ ≫ *η*_*T*_ *> η*_*a*_ *> ϑ*^h^ *> η*_*D*_.

**Figure 7.**
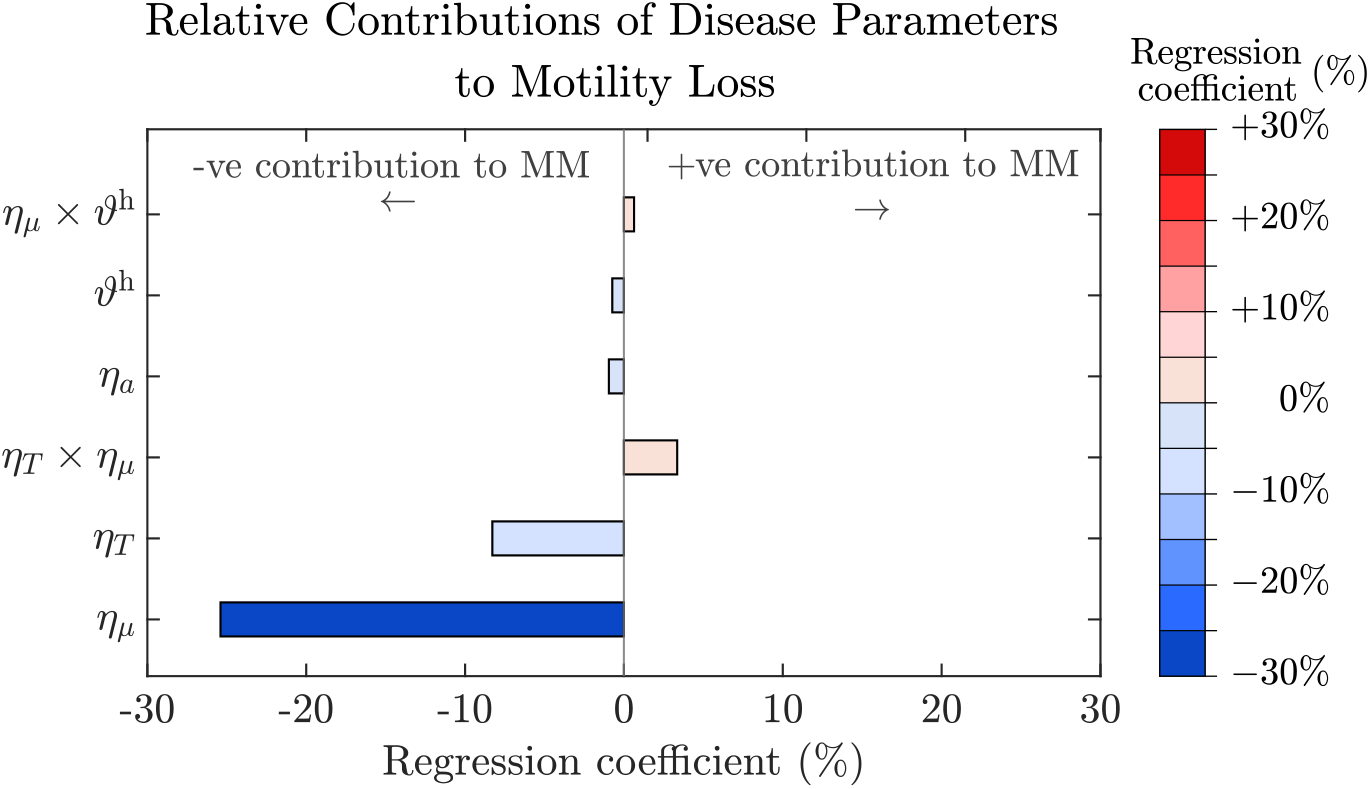
Regression coefficients from the full-factorial 2^5^ model fitted to the 32 ΔMM responses, sorted by absolute magnitude. Terms with |*β*| *<* 0.5 percentage points (pp) are omitted for visual clarity. Negative coefficients indicate reductions in the motility metric. Wall shear modulus (*η*_*µ*_) is the dominant factor (*β*_*µ*_ = *−*25.40 pp; full effect *−*50.79 pp), followed by peak active stress (*η*_*T*_, full effect *−*16.56 pp). Excitation threshold, hypertrophic remodeling, and electrical diffusivity have substantially smaller main effects. The largest interaction is *η*_*T*_ *× η*_*µ*_ (*β*_*Tµ*_ = +3.35 pp; full effect +6.71 pp).

**Figure 8.**
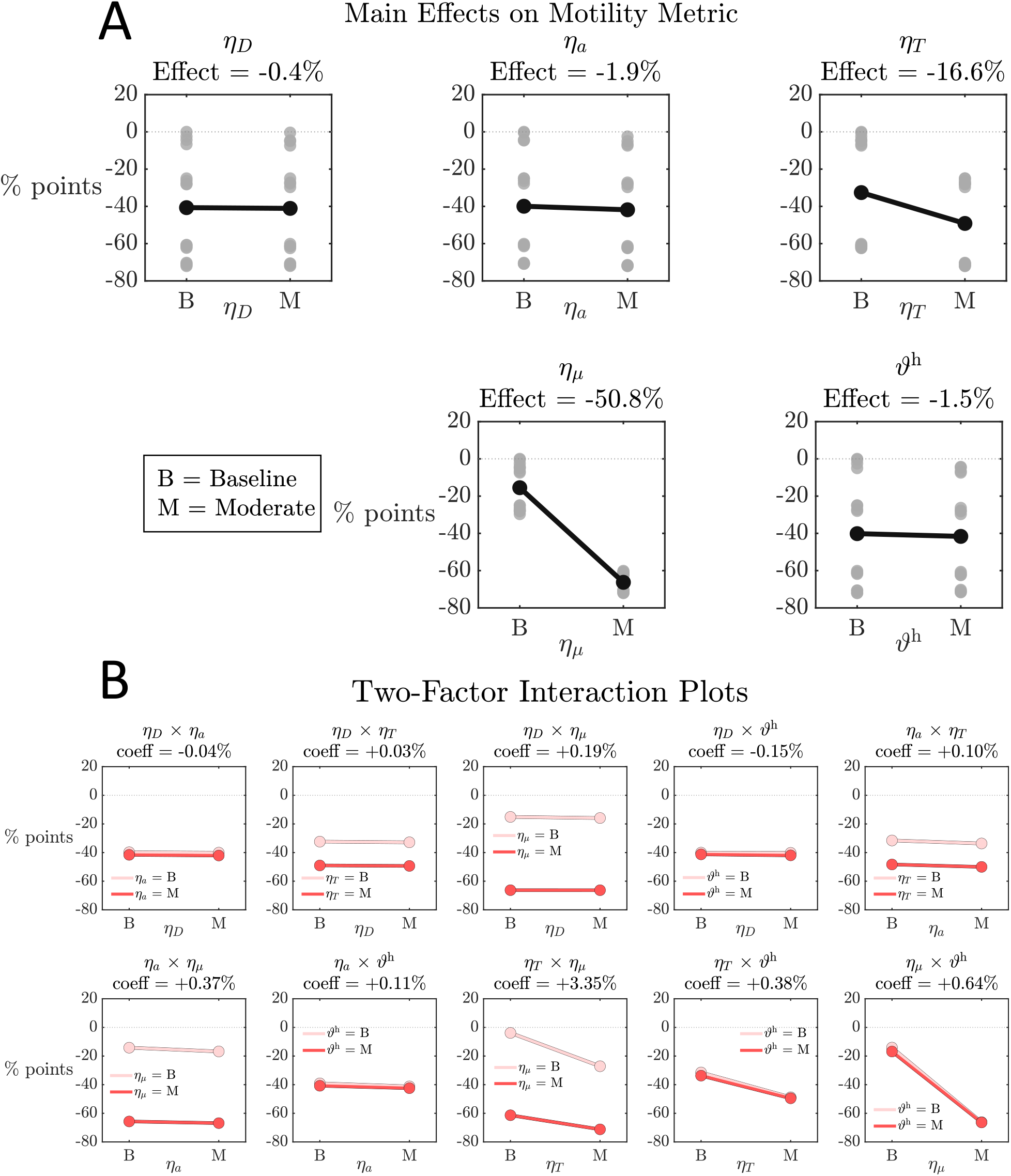
Main-effect and two-factor interaction plots for the 2^5^ DOE. **(A)** Main-effect plots for electrical diffusivity, excitation threshold, peak active stress, wall shear modulus, and hypertrophic remodeling. Wall stiffness produces the largest reduction in the motility metric (*−*50.79 pp), followed by reduced peak active stress (*−*16.56 pp), excitation-threshold alteration (1.89 pp), hypertrophic remodeling (1.47 pp), and reduced electrical diffusivity (*−*0.39 pp). **(B)** Two-factor interaction plots. Pairwise interactions are smaller than the leading main effects, indicating that the response over the tested range is dominated by the main effects.

### 3.7 Two-factor interactions

The two-factor interactions remained smaller than the dominant main effects (Figure 8B). The largest interaction was between impaired contractility and wall stiffness, *η*_*T*_ × *η*_*µ*_, with *β*_*Tµ*_ = +3.35 pp and a full interaction effect of +6.71 pp. The next largest interaction was *η*_*µ*_ × *ϑ*^h^, with a full effect of +1.28 pp, while all remaining interaction effects were below 0.8 pp in magnitude. Overall, the interaction-to-main-effect magnitude ratio was 0.075, indicating that the response over the tested range was dominated by the main effects.

## 4 Discussion

### 4.1 Altered passive mechanics as the primary contributor to dysmotility over the moderate-disease range

This study presents, to our knowledge, an integrated electromechanical finite-element framework for fibrostenosing Crohn’s disease that allows electrical, contractile, mechanical, and structural remodeling mechanisms to be compared within a common computational setting. Across the parameter ranges examined, increased wall stiffness emerged as the dominant contributor to ileal motility loss, with a full effect of −50.79 pp. This effect was approximately three times larger than that of impaired active contractility and more than an order of magnitude larger than the effects of excitation-threshold alteration, hypertrophic remodeling, or reduced electrical diffusivity (Figure 7; Figure 8A).

The mechanistic interpretation follows from the model structure. The wall shear modulus determines the deformation achieved for a given level of active stress. Increasing *η*_*µ*_ therefore increases resistance to circumferential shortening even when electrical activation and active-stress generation remain present. Contractile weakening through *η*_*T*_ acts on the opposite side of the same mechanical balance by reducing the active stress available to deform the wall. Together, the dominance of these two mechanisms indicates that, over the moderate-disease range examined, motility loss is governed primarily by the conversion of electrical activation into mechanical deformation.

### 4.2 Hypertrophic remodeling has less effect on motility loss

Structural hypertrophy produced a comparatively small negative average main effect, with a full effect of −1.47 pp and an average motility-metric ratio of 0.976 (Figure 8A). Thus, the remodeled geometry modestly reduced cyclic lumen-volume deformation when averaged across the other electrical, contractile, and passive-material factor levels. Its effect was substantially smaller than those of wall stiffening and impaired contractility and slightly smaller than that of excitation-threshold alteration. These results indicate that, over the range examined, geometric remodeling alone contributes less to motility loss than the altered mechanical resistance and contractile impairment associated with fibrostenosing disease. A narrower lumen changes the geometry over which contraction occurs, whereas increased tissue stiffness directly limits the strain generated by active smooth-muscle tension. The interaction between wall stiffness and hypertrophic remodeling, *η*_*µ*_ × *ϑ*^h^, had a full effect of only +1.28 pp, further indicating that these structural mechanisms do not exhibit strong synergistic coupling over the range examined. The present ranking should therefore be interpreted as a comparison among five selected disease-associated mechanisms rather than as an exhaustive ranking of all processes involved in fibrostenosing Crohn’s disease. Hypertrophy itself arises downstream of inflammation and fibrosis, as does the increase in tissue stiffness, and additional structural processes such as submucosal remodeling, altered collagen architecture, stricture length, and patient-specific luminal morphology were not independently varied.

### 4.3 Electrical coupling and the dependence of mechanism ranking on disease severity

Reduced electrical diffusivity produced only a −0.39 percentage-point main effect over the range examined. This is not evidence that electrical coupling is unimportant in fibrostenosing Crohn’s disease; it is evidence that it is unimportant over the range tested. The monodomain relation 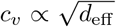 makes the consequence of *η*_*D*_ strongly non-linear. At the moderate level considered here (*η*_*D*_ = 0.46), the effective diffusivity falls to 54 % of its healthy value and the corresponding conduction velocity is expected to fall to approximately 74 % under the monodomain scaling 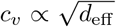 Despite this reduction, the resulting effect on cyclic lumen-volume deformation remained small compared with the mechanical and contractile factors. At the severe-disease value *η*_*D*_ = 0.92, however, the diffusivity falls to 8 % and the conduction velocity to 28 %, a qualitatively different regime in which activation may become spatially localized within the fibrotic segment. More severe obstructive disease may likewise enter a qualitatively different electrophysiological regime. In murine partial small-bowel obstruction, ICC networks were disrupted oral to the obstruction, accompanied by marked attenuation or loss of electrical slow waves and enteric neural responses (Chang et al. 2001). The present moderate-disease DOE does not attempt to reproduce this conduction-failure regime. The effect of excitation-threshold reduction via *η*_*a*_ was similarly small (−1.89 pp), consistent with both electrical parameters acting on activation (Figure 8A).

### 4.4 Comparison with prior models and literature measurements

Methodologically, the framework combines three established modeling traditions. First, the growth step follows the multiplicative decomposition of Rodriguez et al. (1994), with radial growth driven by a Fisher-KPP scalar field, applied here to Crohn’s strictures as a 3D continuum model for the first time. Secondly, the electromechanical step builds on the FitzHugh-Nagumo monodomain formulation, widely used in excitable-tissue modeling (Aliev et al. 2000; Clayton et al. 2011). This approach has been adapted to gastric and intestinal smooth muscle (Lin et al. 2006b; Klemm et al. 2020; Klemm et al. 2023) and to colonic intestinal motility (Djoumessi et al. 2024; Djoumessi et al. 2025), but none of these prior intestinal models has incorporated disease-associated changes in electrical diffusivity, excitation threshold, active contractilit, and stiffness simultaneously. Finally, the diffusivity scaling *η*_*D*_ is motivated by the continuous-cable relationship 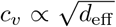 (Kléber and Rudy 2004; De Bakker et al. 1993) and provides a practical surrogate for reduced electrical coupling in the absence of direct intestinal measurements under graded fibrosis.

Our ranking of the impact of each factor may be contrasted with the analogous analysis in cardiac tissue by Telle et al. (2025), where they applied a fractional factorial design to nine fibrosis-associated parameters in patient-specific left-atrial electromechanical models and found impairment of the L-type calcium current to be the dominant determinant of atrial function, with conduction velocity and stiffness altering their metrics by less than 2 %. Contractile impairment is influential in both studies, but unlike the cardiac analysis, our model identifies passive stiffness as the dominant determinant of motility loss. Their stiffness perturbation was calibrated to a two-fold change in load at 5 % stretch and applied within a thin-walled chamber, whereas our *η*_*µ*_ raises the shear modulus by 165 % in a thick-walled tube whose function depends on near-complete luminal occlusion. Occluding a lumen requires far larger strains than reducing a chamber volume by a third, so a given increase in stiffness costs more.

Our predicted reductions in motility are broadly consistent with clinical observations: Menys et al. (2013b) reported motility scores of 0.43 au in normal bowel versus 0.15 au in strictured bowel (a reduction of approximately 65 %) in 91 patients undergoing MR enterography. With all five disease-associated factors at their moderate-disease levels, the model predicts ΔMM = −71.6% in Run 32, which is similar in magnitude to the approximately 65 % reduction reported clinically. Beek et al. (2025) found no significant difference in motility between inflammatory and chronic fibrotic strictures (*p* = 0.6); our finding that wall stiffness, impaired active contractility, and hypertrophic remodeling dominate the response, while the electrical parameters contribute comparatively little over the tested range offers one possible explanation, in that the mechanical consequences of a stricture are largely shared between the two histopathological subtypes.

### 4.5 Clinical and therapeutic implications for fibrostenosing Crohn’s disease

Altered passive mechanics produced the largest motility reduction over the range examined (Figure 7), suggesting that preventing or limiting wall stiffening may be important for preserving mechanical function in fibrostenosing Crohn’s disease. The finding that hypertrophic remodeling also modestly reduced motility indicates that luminal geometry and passive material properties contribute through distinct mechanisms. No antifibrotic therapy is currently approved for Crohn’s disease (Rieder et al. 2018; Bettenworth et al. 2024), and current anti-inflammatory agents and biologics do not reverse muscularis hypertrophy or wall stiffening. Endoscopic balloon dilation restores luminal caliber in approximately 90 % of procedures but is followed by symptomatic recurrence in roughly 70 % of patients and re-intervention in approximately one third (Bettenworth et al. 2017). Our results might offer a mechanical rationale for reduced motility: dilation enlarges the lumen but leaves wall stiffness unchanged, and this residual stiffness continues to limit contraction in the model. The same mechanical relationship suggests that stiffness measured by MR or ultrasound elastography may carry more prognostic information about motility than wall thickness. These predictions require experimental and clinical validation and are conditional on the moderate-disease range over which the parameters were varied. One potential readout for testing this predicted relationship between wall mechanics and motility, as well as the effects of future interventions, is cine-MR enterography quantification of small-bowel motility, already validated against symptoms, endoscopy, and biologic response (Menys et al. 2018; Plumb et al. 2025; Steiner et al. 2022).

### 4.6 Limitations and future work

Several limitations of the present framework should be acknowledged. The fibrosis field *m* is represented phenomenologically using a Fisher-KPP equation rather than through explicit inflammatory and fibrotic pathways involving TGF-*β*, myofibroblasts, and creeping fat (D’Alessio et al. 2022; Liu et al. 2025), while the four Model 2 disease-scaling parameters *η*_*D*_, *η*_*a*_, *η*_*T*_, and *η*_*µ*_ represent selected functional consequences of inflammation and fibrosis. Hypertrophic remodeling is treated separately through the Model 1 growth multiplier *ϑ*^h^ and the resulting change in geometry. The geometry is also simplified, with radial growth applied to an idealized quarter-cylinder representing the muscularis propria without explicit submucosal remodeling, fiber dispersion, mesenteric tethering, or patient-specific stricture morphology. The FitzHugh–Nagumo formulation provides a reduced description of intestinal electrophysiology and does not resolve cell-specific ionic mechanisms (Corrias and Buist 2007; Poh et al. 2012). The present framework also excludes luminal contents and enteric-nervous-system-mediated neuromechanical feedback and therefore quantifies cyclic wall deformation rather than propulsive efficiency. Colonic propulsion depends strongly on luminal load and enteric neural activity (Costa et al. 2015), while Piezo1-mediated mechanosensation in enteric neurons has recently been implicated in gastrointestinal motility (Xie et al. 2025). Because the present model represents the terminal ileum, these results should not be extrapolated directly to colonic propulsion, where luminal-content and enteric-neural feedback are central components of motor function. Several parameter changes are not directly calibrated from intestinal measurements, particularly *η*_*D*_, because conduction velocity has not been quantified across graded intestinal fibrosis (Kléber and Rudy 2004; De Bakker et al. 1993). The DOE evaluates five selected disease-associated mechanisms and therefore does not constitute an exhaustive sensitivity analysis of fibrostenosing Crohn’s disease. The structural response of a stricture also depends on factors not independently varied here, including fibrotic-segment length, submucosal remodeling, collagen architecture and dispersion, circumferential versus longitudinal muscle remodeling, mesenteric tethering, and patient-specific stricture morphology. Likewise, the enteric nervous system, immune-mediated modulation of ICC and smooth muscle, and more detailed ionic mechanisms are not explicitly represented. The resulting ranking is also conditional on the selected perturbation ranges. The physical magnitude of the B-to-M change differs among factors: for example, the moderate stiffness level corresponds to *η*_*µ*_ = 1.655, whereas *η*_*a*_ and *η*_*T*_ are both varied to 0.25. Consequently, the dominance of wall stiffness reflects both the sensitivity of the model to mechanical resistance and the magnitude of the imposed disease-associated perturbation. More severe electrical dysfunction, including regimes approaching conduction block, was not examined.

Future work will move toward patient-specific geometries reconstructed from MR enterography, calibration of the diffusivity and stiffness fields against MR elastography, cine-MRI and high-resolution manometry based motility measurements (Wilkens et al. 2022; Plumb et al. 2025; Anantha Krishnan et al. 2026a), coupling to enteric nervous system and immune compartments, replacement of FitzHugh-Nagumo with biophysical ICC-SMC ionic models (Lin et al. 2006a; Djoumessi et al. 2025), and *in silico* screening of candidate anti-fibrotic interventions to prioritize targets before clinical translation.

## 5 Conclusions

Our integrated electromechanical framework identifies altered tissue mechanics as the primary contributor to motility loss in fibrostenosing Crohn’s disease over the parameter ranges examined. In the full-factorial 2^5^ analysis, increased wall stiffness produced the largest reduction in cyclic lumen-volume deformation, followed by impaired smooth-muscle contractility, whereas excitation-threshold alteration, hypertrophic remodeling, and reduced electrical diffusivity had substantially smaller effects. Pairwise interactions were smaller than the dominant main effects, indicating that the response over the tested range was governed primarily by the individual main effects. These findings indicate that passive wall resistance, active force generation, and structural remodeling make distinct contributions to fibrostenosis-associated dysmotility and highlight the mechanical conversion of activation into lumen deformation as a key determinant of ileal function.

## Acknowledgments

A.A.K. acknowledges helpful discussions with Dr. Alex Menys (Motilent Limited, UK) regarding cine-MRI measurements. A.A.K. acknowledges Dr. Nick Spencer (Flinders University) for his valuable feedback on the project.

## Contributions

Conceptualization: A.A.K.

Methodology: A.A.K., M.A.H.

Software: A.A.K.

Formal analysis: A.A.K., M.A.

Investigation: A.A.K.

Writing: A.A.K., M.A.

Writing - Review & Editing: A.A.K., M.A., M.A.H.

Supervision: M.A.H.

Project administration: M.A.H.

Funding acquisition: M.A.H.

## Data availability

The model files are publicly available at: https://github.com/theCoMMaNDlab/Crohns_motility.

